# *RUNX1* aberrations in blast-phase CML induce the RBP SPATS2L which promotes growth, survival and stress granule assembly

**DOI:** 10.64898/2026.03.27.709496

**Authors:** DA Palmer, AL Muir, R Craig, PA Lewis, M Wilson, K Heesom, GA Horne, M Copland, S Mustjoki, S Adnan-Awad, K Porkka, S Jain, N Bayat, D Famili, H Webb, MJ West, FK Mardakheh, T Chevassut, A Tonks, SG Kellaway, BP Towler, RG Morgan

## Abstract

The RUNX1 transcription factor is a critical regulator of hematopoiesis and frequently mutated in myeloid malignancies. In the myeloproliferative neoplasm, chronic myeloid leukemia (CML), secondary somatic *RUNX1* mutations and *RUNX1::MECOM/EVI1*, are associated with tyrosine kinase inhibitor (TKI) resistance and progression to the blast-phase (BP-CML). Research has predominantly focussed on transcriptional dysregulation mediated by *RUNX1* mutations in myeloid malignancies, whilst post-transcriptional dysregulation remains comparatively unexplored. To address this, we used orthogonal organic phase separation (OOPS), to characterise the RNA-binding proteome of RUNX1 deficient BP-CML cells. RUNX1 depleted BP-CML cells exhibited significant alterations to RBP abundance involved in stress response pathways and translation/ribosome-biogenesis (RiBi). Furthermore, RUNX1 depletion or expression of RUNX1::EVI1 in BP-CML cells induced expression and RNA binding activity of SPATS2L, a component of stress granules (SG); membraneless cytoplasmic condensates protecting mRNAs from degradation, promoting survival under stress. Whilst RUNX1 depletion increased SG-assembly, SPATS2L depletion reduced SG-assembly in BP-CML cells and inhibited the growth and survival of multiple BP-CML cell lines. The translation inhibitor homoharringtonine (HHT), used historically in TKI-resistant CML, ablated SG-assembly in BP-CML cells with RUNX1 depletion, and, primary BP-CML cells with LOF/hypomorphic *RUNX1* mutations (characterised by defective DNA-binding/CBFβ-interaction) were preferentially sensitised to HHT. Finally, suppressing SPATS2L expression induced by RUNX1 depletion, increased the HHT-sensitivity of RUNX1 depleted BP-CML cells, suggesting SPATS2L contributes to therapeutic resistance in CML with *RUNX1* mutations. This study suggests that SPATS2L and SG induction could be critical to *RUNX1*-mutant leukemias, and, provides preliminary evidence for a mutationally-targeted approach in CML with *RUNX1* aberrations.

## Introduction

The myeloproliferative neoplasm, chronic myeloid leukemia (CML), is characterised by the *BCR::ABL1* mutation, which drives clonal proliferation and genomic instability.^1^ Tyrosine kinase inhibitors (TKIs), have markedly improved survival-rates in chronic-phase CML (CP-CML).^2–8^ However, in blast-phase CML (BP-CML), responses to single-agent TKIs are often short-lived, with a median-response duration across all TKIs of less than a year.^9^ This necessitates the use of intensive chemotherapy consolidated with an allogeneic stem cell transplant to induce durable remissions, presenting significant toxicities.^10^ The acquisition of secondary somatic mutations in CML, seen in the driver-genes associated with myeloid malignancies, or, additional chromosomal abnormalities (ACAs), accelerates transformation into the advanced-phases.^11–14^ Somatic *RUNX1* mutations are enriched in BP-CML, occurring at a rate of 10-30%,^11,12^ but are seldom detected at diagnosis,^13,15^ indicating they serve as critical molecular events driving TKI resistance and disease transformation. Furthermore, the t(3;21)(q26.2;q22) translocation, generating *RUNX1::MECOM/EVI1* (MDS1 and EVI1 complex locus) transcripts, encountered in BP-CML as part of the high-risk 3q.26 ACA-group,^14^ is also associated with TKI resistance and adverse survival.^14,16^

Myeloid malignancies are frequently enriched for *RUNX1* mutations and *RUNX1* fusion genes.^17,18^ *RUNX1* mutations are associated with chemoresistance and adverse survival, occurring as missense, frameshift(indels), nonsense and splice site mutations, frequently clustering in the DNA-binding, runt homology domain (RHD).^11–13, 19–23^ However, in CML, somatic *RUNX1* mutations have also been shown to cooperate with *BCR::ABL1,* accelerating leukaemogenesis.^24^ Furthermore, *RUNX1* mutations in BP-CML, are associated with the induction of interferon signalling, immune-pathways and lymphoid antigen expression,^23^ and pre-clinical responses to novel therapies including mTOR, BCL2 and VEGF inhibitors has been shown.^23^ This indicates *RUNX1* mutations in CML are associated with their own unique mechanisms for disease transformation, which could be amenable to targeting. *RUNX1::MECOM/EVI1* disrupts *RUNX1-* and *EVI1-* driven gene networks, affecting hematopoietic differentiation and lineage commitment.^25–29^ *RUNX1::MECOM/EVI1* has also been shown to cooperate with *BCR::ABL1,* inducing leukaemic progression.^30^ Consequently, targeting *RUNX1* aberrations in CML remains an area of unmet clinical need.

Research has predominantly focussed on transcriptional dysregulation mediated by *RUNX1* mutations or fusion genes in myeloid malignancies.^17,18^ Here, we characterised changes to the RNA-bound proteome in BP-CML cells with RUNX1 depletion. Stress response pathways and ribosome biogenesis (RiBi)/translation were enriched processes for differentially-regulated RNA-binding proteins (RBP) upon RUNX1 depletion. Furthermore, RUNX1 depletion or expression of RUNX1::EVI1, induced the RBP, SPATS2L (Spermatogenesis Associated Serine Rich 2-Like or Stress Granule and Nucleolar Protein), previously identified as upregulated in *RUNX1* mutated primary BP-CML cells.^23^ SPATS2L is a component of stress granules (SG);^31,32^ membraneless organelles which promote survival and therapeutic resistance.^33,34^ We found SPATS2L depletion inhibited growth, survival and SG-assembly in BP-CML cells. Furthermore, BP-CML cells with RUNX1 depletion had increased SGs, which were ablated with the translation-inhibitor homoharringtonine (HHT). Finally, primary *RUNX1* mutated BP-CML cells with LOF/hypomorphic *RUNX1* mutations were preferentially sensitised to HHT. Overall, these findings support a mutationally-targeted approach in CML with *RUNX1* aberrations and demonstrate that disrupting SG-assembly could represent a novel therapeutic strategy.

## Methods

### Orthogonal Organic Phase Separation (OOPS)

OOPS was performed as previously described,^35^ adapted to suspension cells. For each experimental replicate ∼10×10^6^ cells were crosslinked with 300mJ of UV-C then pelleted before resuspension in 1mL of TRIzol™ LS Reagent (Invitrogen, ThermoFisher Scientific) followed by the addition of 200μL of UltraPure™ Chloroform (Invitrogen, ThermoFisher Scientific). Following vortexing and centrifugation at 4°C, 12,000 x *g*, for 15 minutes, the aqueous (clear-layer) and organic phase (pink-layer) were removed, isolating the interphase (white-layer) into 1mL of TRIzol™ LS with 200μL UltraPure™ Chloroform (this process was repeated three times). The final interphase was centrifuged for 2 minutes at 4°C, 12,000 x *g*, after the addition of 1mL of ice-cold methanol (ThermoFisher Scientific). Samples were resuspended in 100μL of RBP buffer containing 100mM TEAB (Triethylammonium bicarbonate buffer; Merck-Millipore) 1mM MgCl2 (ThermoFisher Scientific), 1% SDS solution (Severn Biotech) diluted in a stock of RNAase free water (ThermoFisher Scientific), and RNA digestion was carried out with 8μg of RNA-ase A/T1 enzyme (ThermoFisher Scientific). The next day TRIzol™ LS and UltraPure™ Chloroform was added followed by centrifugation (as above), and 400μL of the organic phase was added in a 1:4 ratio to 99+% Acetone (ThermoFisher Scientific) and precipitated overnight at −20°C. Samples were centrifuged at 17,000 x *g*, 4°C for 20 minutes, and resuspended in 50μL RIPA buffer (ThermoFisher Scientific). Protein was quantified in each replicate using an EZQ protein quantitation kit (ThermoFisher Scientific) and 10-15 experimental replicates for each cell line condition were combined into three replicates, each containing 20μg of protein for tandem-mass-tag (TMT)-labelling and LC-MS (described in supplementary methods).

### Other methods

Cell lines/culture, western blotting, lentiviral production/transduction, RNA extraction/purification, quantitative reverse transcription polymerase chain reaction (qRT-PCR), polysome profiling, flow cytometric analysis (7AAD and Annexin V staining), cloning, nuclear/cytoplasmic fractionations, immunofluorescence/confocal laser scanning microscopy (CLSM) and preparation/analysis of primary BP-CML samples is described in the supplementary methods.

## Results

### Induction of stress response pathways and dysregulated RiBi/translation is a feature of BP-CML cells with RUNX1 depletion

To first characterise whether loss of RUNX1 modified the RNA-bound proteome, we used two unique shRNAs targeting the RUNX1 NM_001754.4 transcript (isoform c) in KU812 and K562 myeloid BP-CML cells (Figure 1A), followed by OOPS (a method of enriching RNA-protein interactions) coupled to TMT-labelled LC-MS. We first confirmed RBP enrichment in these OOPS samples using the GO search term ‘RNA-binding’ (*Supplemental Figure S1A-B*). In KU812 cells, 19.2% of RBPs detected (136/708) were altered in abundance with RUNX1 depletion compared to controls (p<0.05; Figure 1B), whilst fewer changes were observed in K562 cells (8/963); thus, we focussed our downstream analysis on KU812 cells. To establish pathways enriched for differentially-regulated RBPs, we performed an analysis of canonical pathways (Figure 1C-D). RUNX1 depletion activated multiple pathways associated with the stress response, such as, *response of EIF2AK4 (GCN2) to amino acid deficiency*, *oxidative phosphorylation*, *NRF2−mediated oxidative stress response*, and the *unfolded protein response* (UPR). Furthermore, there was an enrichment of differentially-regulated RBPs associated with RiBi/translation, including with, *rRNA processing in the nucleolus and cytosol* and *translation-initiation* -*elongation* and *-termination*.

**Figure 1.**
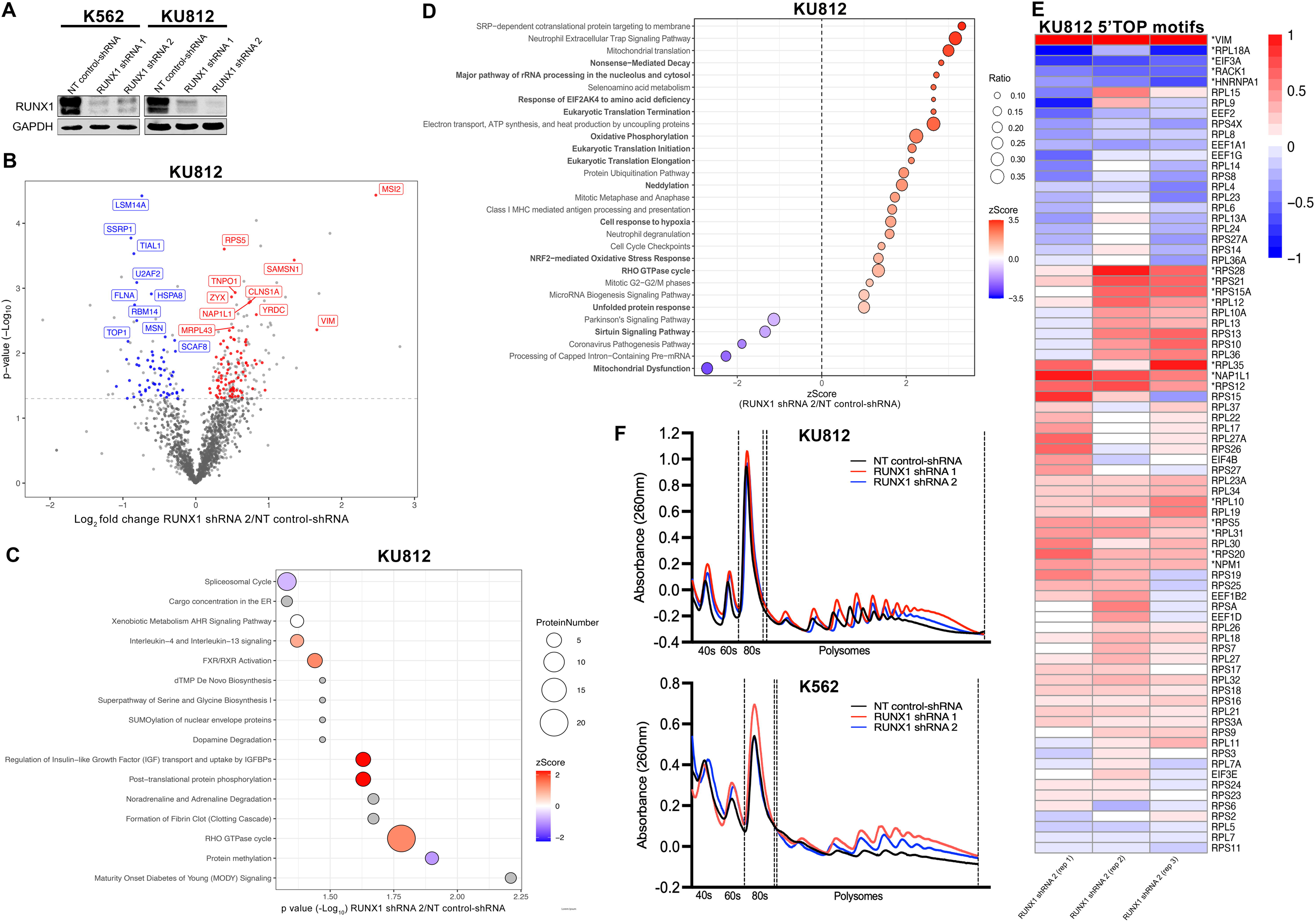
RUNX1 depleted BP-CML cells are enriched for differentially regulated RBPs associated with stress response pathways and dysregulated RiBi/translation. A) Western blots of K562 and KU812 cells transduced with two unique RUNX1-targeting shRNAs compared to a non-targeting (NT) control-shRNA. GAPDH was used as a loading control. B) Volcano plot depicting the most differentially-regulated RBPs in RUNX1 depleted (shRNA 2) (*n = 3*) versus NT control-shRNA KU812 cells (*n = 3*). The most significantly enriched (red)/depleted (blue) RBPs are annotated. Horizontal dashed lines indicate the threshold for statistical significance (p<0.05) on a -log_10_ scale (1.3). Dark grey dots are non-statistically significant RBPs and light grey dots proteins which are non-RNA-binding. C) A reactome of canonical pathways enriched for differentially regulated-RBPs in RUNX1 depleted (shRNA 2) versus NT control-shRNA KU812 cells. Pathways coloured in grey represent insufficient proteins significantly changed, with known activation/inhibition roles within that pathway to calculate a z-Score. D) A reactome of canonical pathways enriched for differentially regulated-RBPs in RUNX1 depleted (shRNA 2) versus NT control-shRNA KU812 cells based on z-Scores alone, comparing the direction of LogFC and its alignment, to the expected direction of change, based upon activation/enrichment of that pathway. A z-Score >-1 and <1 was used as a cut-off, and pathways represented by ≤5 proteins were excluded. Relevant pathways have been highlighted. E) A heat map containing all RBPs from OOPS in RUNX1 depleted (shRNA 2) KU812 cells with 5’TOP motifs. Each RUNX1 shRNA replicate is compared to its respective NT control-shRNA replicate (*n = 3*). LogFC is represented by the colour scale. Statistically significant proteins are annotated by *p<0.05. F) Polysome profiles of KU812 and K562 cells transduced with NT control-shRNAs (black) compared to RUNX1-targeting shRNAs (red/blue). Dashed vertical lines represent monosome and polysome fractions. The 40s, 60s, 80s subunits, polysomes and absorbance readings (260nm) are annotated.

RUNX1 deficient HSCs have been shown to have attenuated ER stress, an UPR and low-rates of RiBi.^36^ The enrichment of differentially-regulated RBPs associated with stress response pathways and dysregulated RiBi/translation, we found in KU812 cells with RUNX1 depletion, led us to consider whether RUNX1 regulates 5’TOP (terminal oligopyrimidine) motifs. Cells exposed to stress stall translation by downregulating the expression of mRNAs containing a cis-regulatory RNA-element (CRE), known as the 5’TOP motif; a conserved region in the 5’UTR of mRNAs of ribosomal subunit-proteins and components of translational complexes.^37,38^ RUNX1 depletion in KU812 cells led to a significant difference (p<0.05) in the expression of 22% (17/78) of 5’TOP motifs (Figure 1E). Of these differentially expressed proteins, 82% were increased, 65% of which were components of ribosomal subunits. To expand these findings, we performed polysome profiling in KU812 and K562 cells with RUNX1 depletion to evaluate ribosome activity (Figure 1F; *Supplemental Figure S2*). Area under the curve (AUC) analysis showed an increase in monosomes and polysomes with RUNX1 depletion, but no increase in the polysome/monosome ratio (except in K562 RUNX1 shRNA 2 cells where it was increased). Overall, these findings indicate RUNX1 depletion increases ribosome abundance in polysomes in BP-CML cells, without causing a shift towards an increase in global translation.

### RUNX1 depletion or expression of RUNX1::EVI1 causes induction of SPATS2L in BP-CML cells

We next decided to interrogate an RBP with dysregulated RNA-binding activity/expression in the post-transcriptional processes identified, common to both K562 and KU812 cells. We identified the upregulation of SPATS2L (Figure 2A-B), an RBP which regulates rRNA processing,^31^ found to be enriched 1.65 and 0.52 log_2_ fold bound to RNA derived from K562 and KU812 cells upon RUNX1 depletion, respectively. We subsequently observed increased expression of multiple SPATS2L isoforms from whole cell lysates derived from both cell lines with RUNX1 depletion (Figure 2C), indicating total expression was also impacted. To consider other forms of RUNX1 dysregulation on SPATS2L expression in the context of BP-CML, we evaluated whether expression of *RUNX1::EVI1* t(3;21), part of the 3q.26 cytogenetically-rearranged group,^14^ which has a repressive effect over *RUNX1*,^25–29^ could also induce SPATS2L. To do this, we induced expression of *RUNX1::EVI1* via lentiviral transduction of K562 and KU812 cells. RUNX1 and EVI1 appeared between 150-250 kDa, confirming that the N-terminus of RUNX1 was fused to EVI1 (Figure 2D). We found RUNX1::EVI1 induced SPATS2L expression in each of these cells, comparable to endogenous levels present in SKH1 cells; a BP-CML cell line harbouring t(3;21)^39^ (Figure 2D). Given, previous reports have shown *SPATS2L* is induced in *RUNX1* mutated versus wild-type primary BP-CML cells,^23^ we next considered whether SPATS2L induction could be derived from elevated transcript levels. We demonstrated a significant increase in *SPATS2L* mRNA expression with both expression of RUNX1::EVI1 and RUNX1 depletion (Figure 2E-F). Taken together, these findings demonstrate SPATS2L is induced in BP-CML cells upon different forms of RUNX1 dysregulation, implying a wider role in BP-CML with *RUNX1* aberrations.

**Figure 2.**
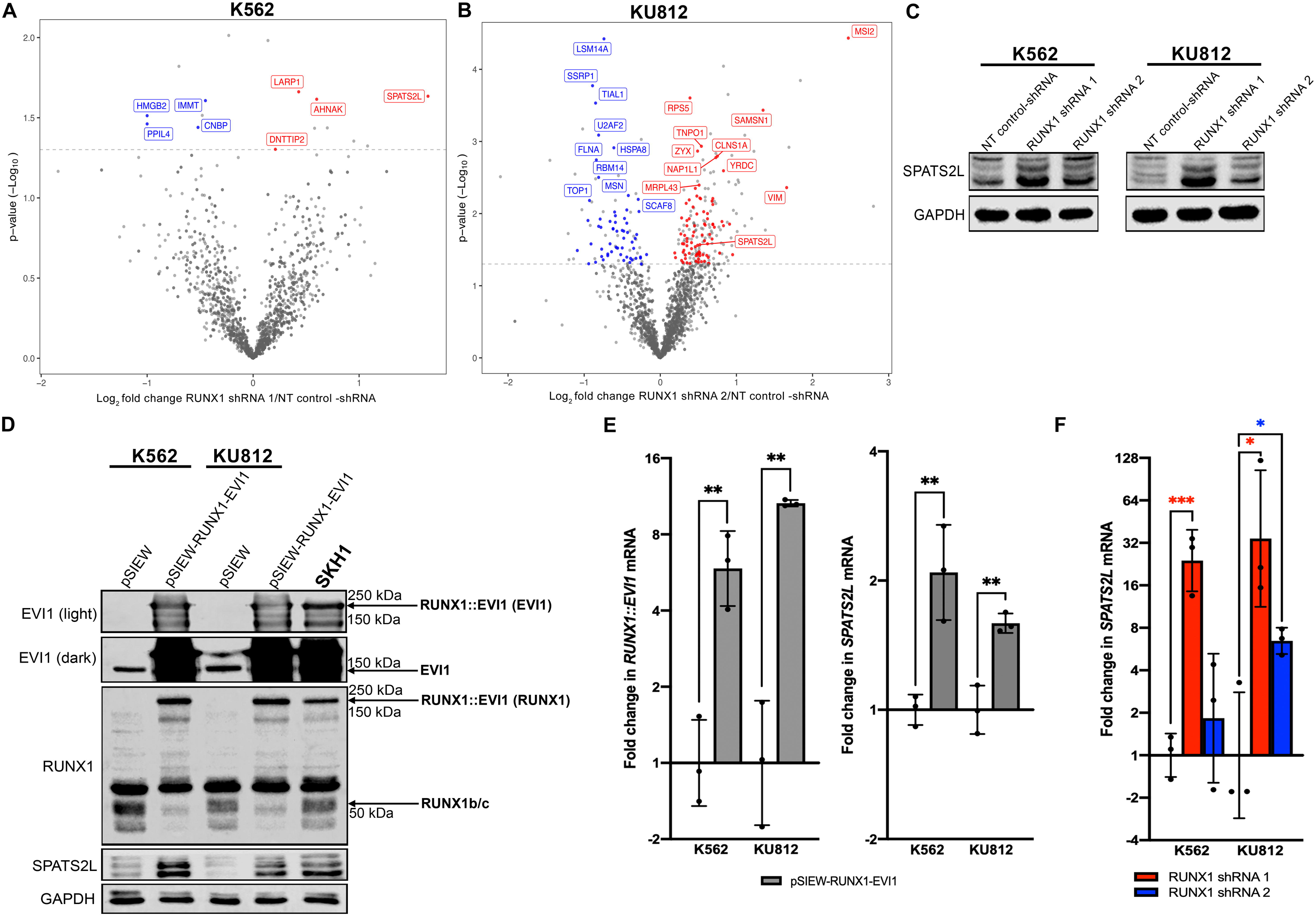
RUNX1 depletion and expression of RUNX1::EVI1 induces SPATS2L in BP-CML cells. **A-B)** Volcano plots of the most differentially-regulated RBPs in RUNX1 depleted K562 (shRNA 1) and KU812 (shRNA 2) cells (*n = 3*) versus NT-control shRNAs (*n = 3*). Each plot depicts the most significantly enriched (red)/depleted (blue) RBPs. Horizontal dashed lines indicate the threshold for statistical significance (p<0.05) on a -log_10_ scale (1.3). SPATS2L has been annotated in each volcano plot. **C)** Western blot showing the level of RUNX1 and SPATS2L in K562 and KU812 cells with RUNX1 depletion compared to NT controls. GAPDH was used as a loading control in each blot. **D)** Western blot showing the level of EVI1, RUNX1 and SPATS2L in K562 and KU812 cells transduced with a pSIEW-RUNX1-EVI1 or pSIEW (control) plasmid, alongside SKH1 cells. The darker EVI1 image allows for visualisation of EVI1 (145 kDa) in pSIEW-K562 and -KU812 cells. GAPDH was used as a loading control. **E-F)** RT-qPCR of *RUNX1::EVI1* and *SPATS2L* in K562 and KU812 cells transduced with pSIEW-RUNX1-EVI1 (grey) compared to pSIEW controls, and, RT-qPCR of *SPATS2L* in K562 and KU812 cells with RUNX1 depletion (red/blue) compared to controls. Data is represented by fold change on ΔΔCt readings. Data represents mean ± 1 s.d (*n = 3*) and statistical significance is denoted as *p<0.05, **p<0.01, or ***p<0.001 from a two-tailed unpaired student’s t-test.

### SPATS2L depletion inhibits growth, survival and SG-assembly in BP-CML cells

SPATS2L regulates the growth and survival of acute myeloid leukemia (AML) cells, and high expression is associated with inferior overall survival outcomes in AML patients.^40^ To explore SPATS2L in the context of BP-CML, we depleted SPATS2L in K562, KU812 and SKH1 cells using two SPATS2L-targeting shRNAs, which targeted multiple SPATS2L isoforms, inferred by depletion of these bands within the predicted molecular weight range (50-75kDa; Figure 3A). SPATS2L depletion inhibited the growth and viability of each of these BP-CML cell lines (Figure 3B-C), further substantiating its pro-leukaemic role. SPATS2L has been shown to localise to ^31^ and coordinate SG-assembly; ^32^ membraneless organelles which protect mRNAs/proteins from degradation and promote survival during conditions of acute stress.^33^ Given these findings, we explored whether the capacity for SG-assembly was disrupted by SPATS2L depletion in BP-CML cells using a 1-hour sodium arsenite treatment, which enhances their formation for visualisation by CLSM (*Supplementary Figure S3*). Using a G3BP Stress Granule Assembly Factor 1 (G3BP1) fluorophore-conjugated antibody (SG marker) we measured SG number/size in K562 and KU812 cells with SPATS2L depletion (Figure 4A-B). This demonstrated a reduction in the average number of SGs per cell, but with no reduction in size. Representative images of SGs in K562 and KU812 cells with SPATS2L depletion are shown (Figure 4C-D). Overall, these results demonstrate that SPATS2L depletion inhibits growth, viability and SG-assembly in BP-CML cells.

**Figure 3.**
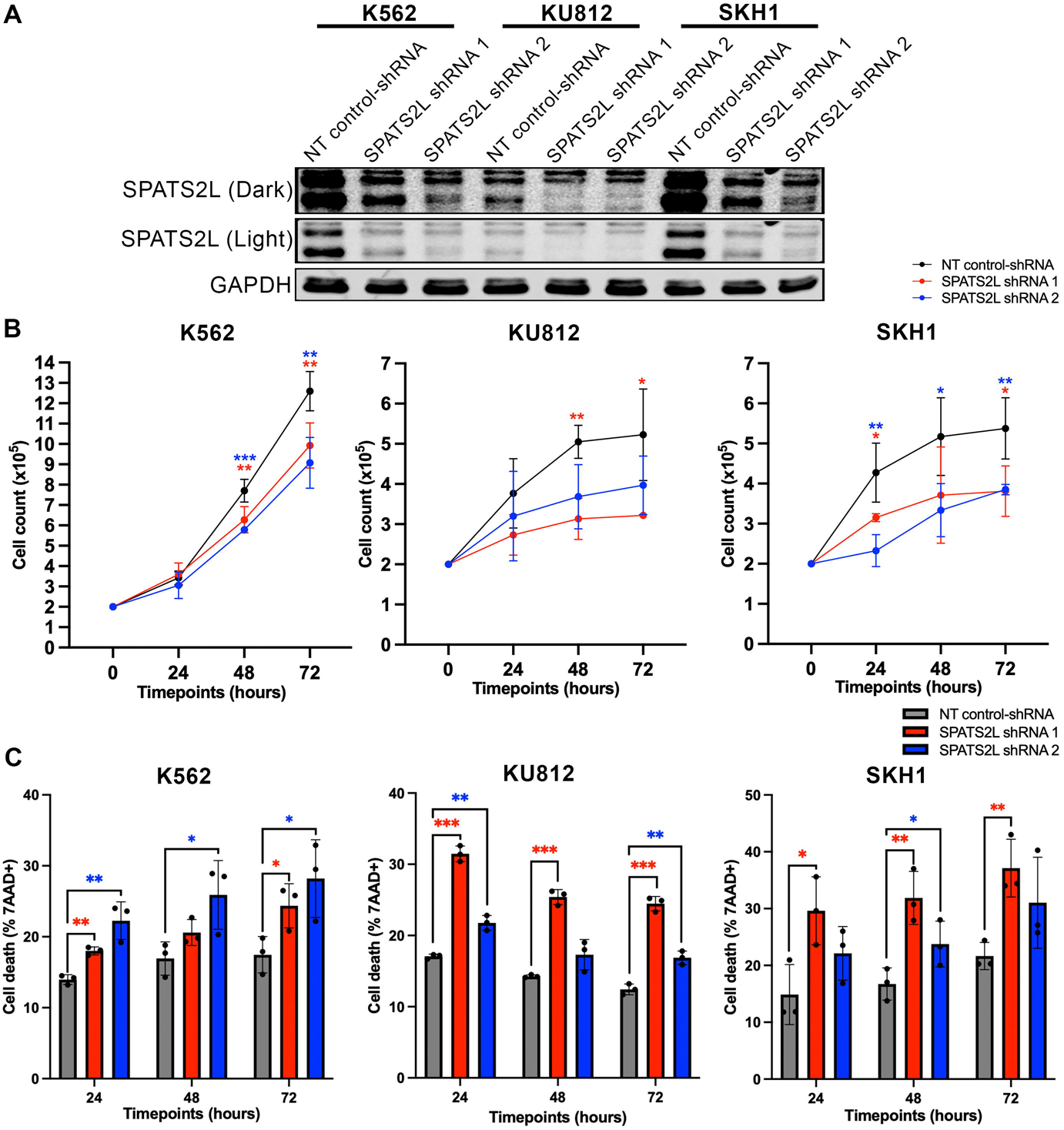
SPATS2L depletion impairs the growth and survival of BP-CML cells. **A)** Western blot showing SPATS2L level in response to two unique SPATS2L-targeting shRNAs in K562, KU812 and SKH1 cells. A dark and light SPATS2L exposure is depicted due to the lower expression in KU812 cells. GAPDH was used as a loading control. **B)** Cell counts in K562, KU812 and SKH1 cells with SPATS2L depletion at 24-48- and 72-hours. Data represents mean ± 1 s.d (*n = 5*) in K562, (*n = 3)* in KU812 and (*n = 4*) in SKH1 cells, with statistical significance denoted by *p<0.05, **p<0.01 or ***p<0.001 from a two-tailed unpaired student’s t-test. **C)** Cell viability in K562, KU812 and SKH1 cells measured by 7AAD^+^ staining (dead cells) at 24-48 and 72-hours. Data represents mean ± 1 s.d (*n = 3*). Statistical significance is denoted by *p<0.05, **p<0.01, or ***p<0.001 from a two-tailed unpaired student’s t-test at each timepoint.

**Figure 4.**
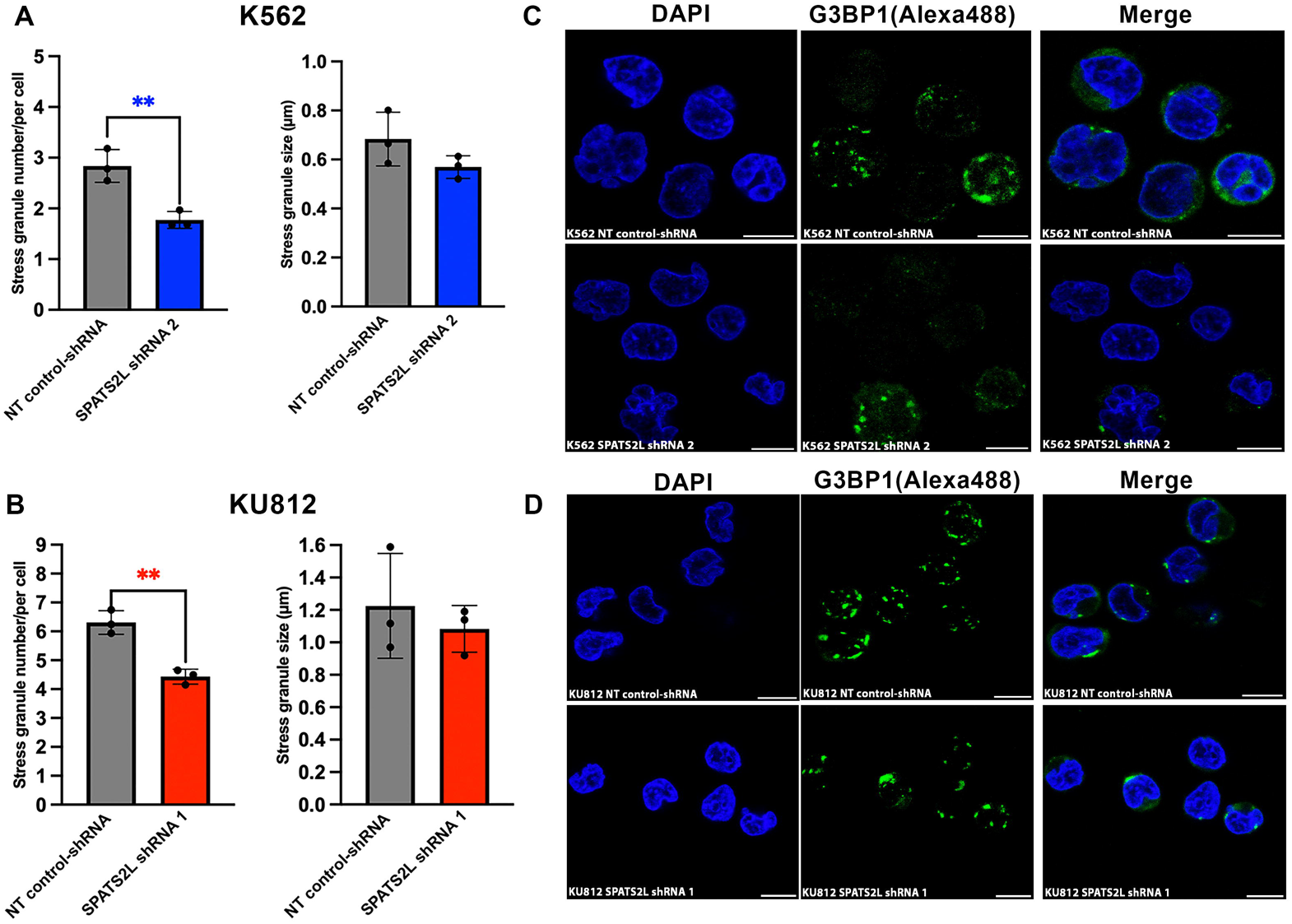
SPATS2L depletion impairs SG-assembly in BP-CML cells. Summary bar graphs showing the average number/size (µm) of SG in SPATS2L-depleted **(A)** K562 and **(B)** KU812 cells relative to NT control-shRNA cells. Data represents mean ± 1 s.d (*n = 3*). Statistical significance is denoted by **p<0.01 from a two-tailed unpaired student’s t-test. Representative CLSM images show the DAPI, G3BP1(Alexa488) and merged stains, of **(C)** K562 and **(D)** KU812 cells transduced with a NT control-shRNA or SPATS2L-targeting shRNAs. White scale bars represent 10µm.

### RUNX1 depletion in BP-CML cells induces SG-assembly which are nearly eradicated with HHT treatment

Upon induction of oxidative stress SPATS2L localises to SGs and the nucleolus, where it has been reported to regulate 5.8s rRNA processing.^31^ Given SPATS2L depletion reduced SG-assembly in BP-CML cells, we evaluated whether the subcellular distribution of SPATS2L was modified by stress induction. To do this we performed nuclear-cytoplasmic fractionations +/− sodium arsenite (SA) treatment in K562 and KU812 cells with RUNX1 depletion. In keeping with our previous observations, we found an overall higher basal level of SPATS2L upon RUNX1 depletion versus controls, which was elevated with SA exposure (Figure 5A). This confirmed SPATS2L activation upon stress induction. Thus, we next considered whether RUNX1 depleted BP-CML cells had increased SG-forming capacity, given our findings that RUNX1 depletion induced multiple RBPs associated with stress response pathways and dysregulated RiBi/translation. We found an increase in SG-assembly in BP-CML cells with RUNX1 depletion, which reached statistical significance (p=0.01) in K562 cells with a similar trend in KU812 cells (p=0.08; Figure 5B-C). Canonical SG-assembly occurs following eIF2α phosphorylation which disrupts cap-dependent translation, leading to the sequestration of mRNAs into these cellular condensates, which lack the 60s ribosomal subunit.^33^ Harringtonine inhibits SA-induced SG-assembly, by freezing 80s ribosomes onto mRNAs at the 60s ribosomal-A site.^41^ HHT has demonstrated *in-vitro* efficacy in AML cells with *RUNX1* mutations,^42^ and we found BP-CML cells with RUNX1 depletion had higher SG-forming capacity following SA exposure. Thus, we hypothesised that an additional mechanism by which HHT induces apoptosis in BP-CML with *RUNX1* aberrations is through SG-inhibition. Supporting this concept, we found SG-assembly was nearly eliminated in K562 RUNX1 shRNA cells following a pre-treatment with HHT and subsequent 1-hour SA exposure (Figure 5D-E). In summary, these data confirm SPATS2L is induced upon stress and RUNX1 depletion increases SG-assembly in BP-CML cells which is prevented with HHT treatment.

**Figure 5.**
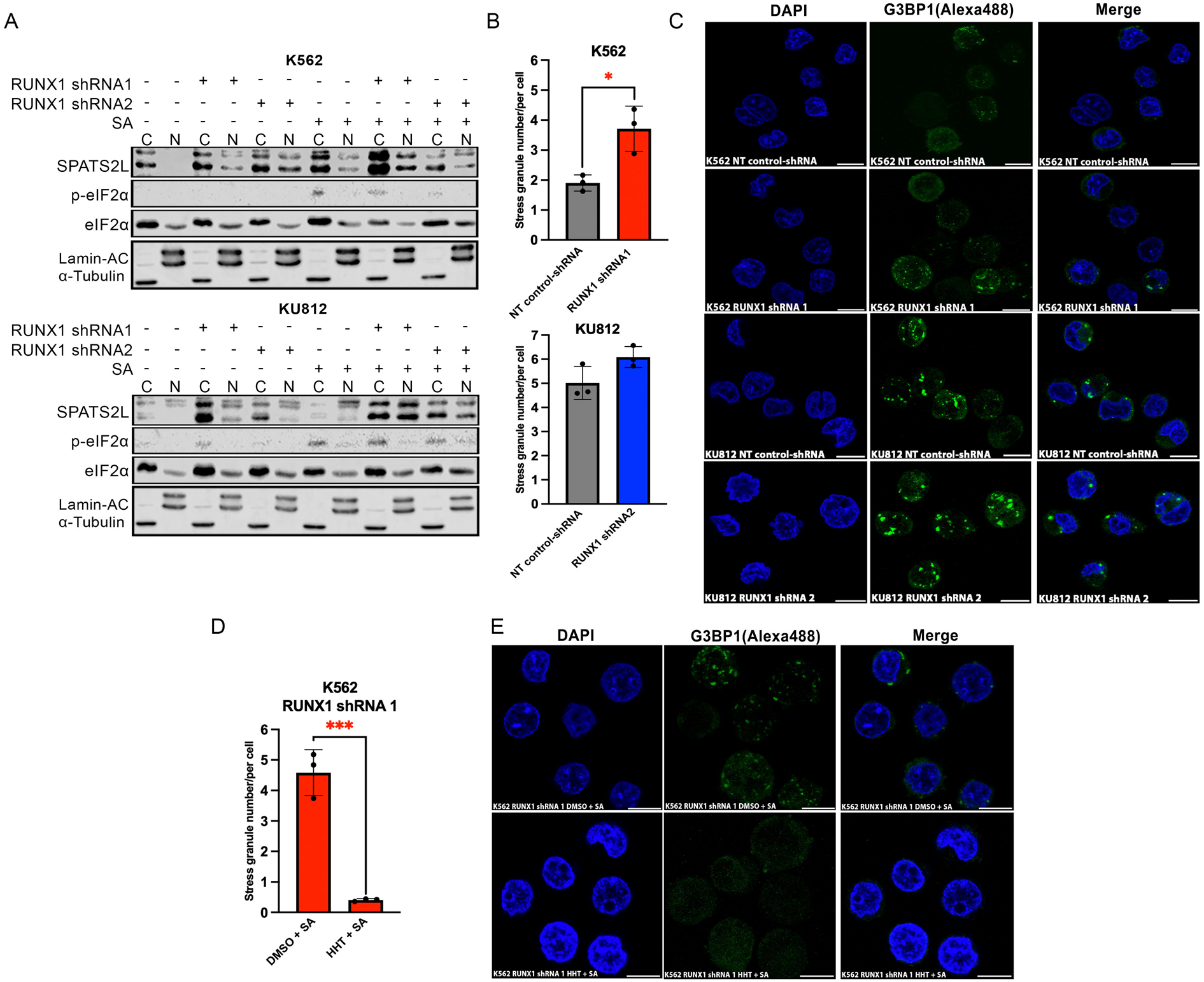
RUNX1 depleted BP-CML cells have increased SG-forming capacity which are eliminated with HHT treatment. **A)** A western blot showing the cytoplasmic (C) and nuclear (N) distribution of SPATS2L, eIF2α and phospho-eIF2α in K562 and KU812 cells with RUNX1 depletion +/− 0.5M sodium arsenite (SA) treatment for 1-hour. Lamin A/C and α-tubulin indicate the level and purity of fractions and p-eIF2α status indicates stress/SG-induction. **B)** Summary graphs showing the average SG number per cell in RUNX1-depleted versus NT control-shRNA K562 and KU812 cells. Data represents mean ± 1 s.d (*n = 3*). Statistical significance is denoted by *p<0.05 from a two-tailed unpaired student’s t-test. **C)** Representative CLSM images showing DAPI, G3BP1 and merged images of K562 and KU812 cells with RUNX1 depletion compared to NT controls. **D)** Summary graph showing the average number of SG per cell in K562 RUNX1 shRNA 1 cells treated with DMSO+SA compared to HHT (50ng/mL for 48-hours)+SA. Data represents mean ± 1 s.d (*n = 3*). Statistical significance is denoted by ***p<0.001 from a two-tailed unpaired student’s t-test. **E)** Representative CLSM images show the DAPI, G3BP1 and merged stains of K562 RUNX1 shRNA 1 cells treated with DMSO+SA compared to HHT+SA. White scale bar represents 10µm.

### BP-CML cells with mutations in RHD of RUNX1 are preferentially sensitised to HHT

Our findings showing HHT nearly eliminated SG-assembly in BP-CML cells with RUNX1 depletion prompted us to further investigate this translation-inhibitor in *RUNX1* mutated BP-CML; a therapy with a history of use in CML.^43–46^ K562 and KU812 cells were sensitised to HHT with RUNX1 depletion, with a heightened-sensitivity after 48-hours (Figure 6A). To explore the mechanism of cell death, we next used the RUNX1 shRNA 1 cell lines, which demonstrated the greatest sensitivity to HHT, to establish whether HHT primarily induced apoptosis at 48-hours (early/late-stage apoptosis measured using Annexin V staining). We found a significant increase in apoptosis with HHT treatment in BP-CML cells with RUNX1 depletion (Figure 6B). However, a higher basal level of apoptosis existed in KU812 RUNX1 shRNA 1 cells compared to controls, therefore we also analysed the sensitivity of these cells using fold change (*Supplemental Figure S4*). Given the increased sensitivity to HHT observed with RUNX1 depletion, we next treated 5 primary *RUNX1* mutated and 4 *RUNX1* wild-type BP-CML samples with HHT (clinical/molecular characteristics of these samples is provided in *Supplementary Table S7).* Overall, we found no significant difference in HHT-sensitivity between *RUNX1*-mutated and wild-type samples (Figure 6C), however, given the heterogeneity of *RUNX1* mutations, we further stratified our analysis based on whether the mutation existed in the RHD or C-terminus. From this, we observed statistically higher rates of HHT-induced apoptosis in BP-CML samples with *RUNX1* RHD mutations versus *RUNX1* wild-type samples (Figure 6D-E). Overall, these findings demonstrate RUNX1 dysregulation sensitises BP-CML cells to HHT through increased apoptosis, with mutations in the RHD demonstrating an increased sensitivity to this treatment.

**Figure 6.**
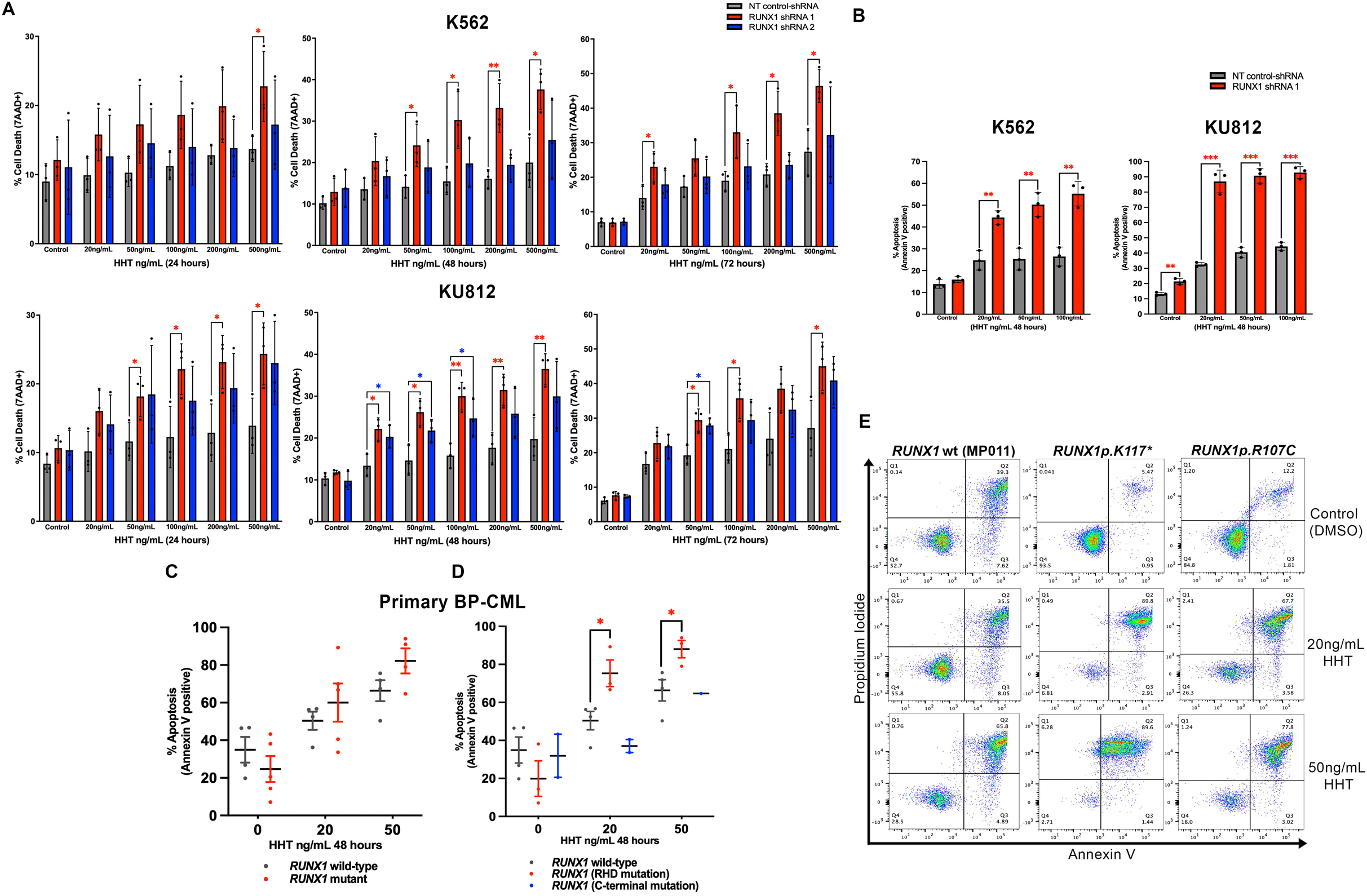
RUNX1 depleted BP-CML cells and primary BP-CML cells with *RUNX1* RHD mutations exhibit enhanced HHT-sensitivity. **A)** Summary bar graphs showing percentage cell death determined by 7AAD positivity in K562 and KU812 cells with RUNX1 depletion compared to NT controls, treated with a dose-response (20-500ng/mL) of HHT for 24- 48- and 72- hours. Data represents mean ± 1 s.d (*n = 3*). Statistical significance is denoted by *p<0.05 or **p<0.01 from a two-tailed unpaired student’s t-test. **B)** Percentage apoptosis (Annexin V positive staining for early/late-stage apoptosis) by way of flow cytometry analysis in KU812 and K562 cells with RUNX1 depletion (shRNA 1) compared to NT controls, treated with 20-100ng/mL HHT at 48-hours. Data represents mean ± 1 s.d (*n = 3*). Statistical significance is denoted by **p<0.01 or ***p<0.001 from a two-tailed unpaired student’s t-test **C)** Percentage apoptosis (early/late-stage) determined by Annexin V positive staining in *RUNX1* mutated compared to *RUNX1* wild-type, patient-derived BP-CML cells, after treatment with 20ng/mL or 50ng/mL HHT at 48-hours (compared to a control dose). Data represents mean ± SEM (*n = 5*) *RUNX1* mutated samples and (*n = 4*) *RUNX1* wild-type samples. One *RUNX1* mutated sample (MP014) had insufficient cells for a 50ng/mL HHT dose. **D)** Percentage apoptosis (early/late-stage) determined by Annexin V positive staining in patient-derived BP-CML cells with/without *RUNX1* mutations, with *RUNX1* mutations grouped based on whether they occur in the RHD (red), or C-terminus (blue), treated with 20ng/mL or 50ng/mL HHT at 48-hours (compared to a control dose). Data represents mean ± SEM (*n = 3*) *RUNX1* RHD mutated samples and (*n = 4*) *RUNX1* wild-type samples. Statistical significance is denoted by *p<0.05 from a two-tailed unpaired student’s t-test. **E)** Representative flow cytometric pseudoplots of Annexin V staining are shown for a *RUNX1* wild-type (MP011) primary BP-CML sample and two primary *RUNX1* mutated BP-CML samples with mutations in the RHD (*RUNX1p.K117\** and *RUNX1p.R107C*), treated with 20ng/mL or 50ng/mL HHT, compared to a control dose.

### SPATS2L inhibition in RUNX1 depleted BP-CML cells inhibits growth, viability and increases HHT sensitivity

SPATS2L amplification occurs with increasing resistance indices to HHT in AML cells and its subsequent depletion sensitises these cells to HHT.^40^ We found dysregulated RiBi/translation and increased SG-assembly was a feature of BP-CML cells with RUNX1 depletion, which could sensitise these cells to HHT. However, SPATS2L could also contribute to therapeutic resistance, by sustaining SG-assembly in response to treatment. Thus, we next aimed to establish whether inhibition of SPATS2L induction observed upon RUNX1 depletion could impact the growth, viability and HHT-sensitivity of BP-CML cells. For this, we created a double-transduced K562 cell line using RUNX1 shRNA 1 (puromycin-selected), coupled with two GFP^+^ SPATS2L-targeting shRNAs. High transduction efficiency was confirmed through GFP^+^ assessment indicating efficient SPATS2L-targeting/control shRNAs in K562 RUNX1 shRNA 1 cells (Figure 7A). We next measured SPATS2L expression, which confirmed the previously observed SPATS2L induction upon RUNX1 depletion was abrogated in the double-transduced K562 cells, most notably using SPATS2L shRNA 2 (Figure 7B). Thus, we used this K562 cell line for subsequent testing, which exhibited overall impaired growth and viability compared with the control K562 RUNX1 shRNA line capable of SPATS2L induction (Figure 7C-D). Finally, to establish whether SPATS2L inhibition could further increase HHT-sensitivity of RUNX1 depleted K562 cells, we treated K562 RUNX1 shRNA1/SPATS2L shRNA 2 cells with a dose response of HHT at 48-hours. Given the higher basal level of death in K562 RUNX1 shRNA1/SPATS2L shRNA 2 cells we measured HHT-sensitivity with fold change (Figure 7E). This demonstrated K562 cells with RUNX1 depletion were further sensitised to HHT when SPATS2L induction is inhibited. Overall, these results support the targeting of SPATS2L in the context of RUNX1 dysregulation in BP-CML, and, could imply SG-assembly may contribute to therapeutic resistance in this setting.

**Figure 7.**
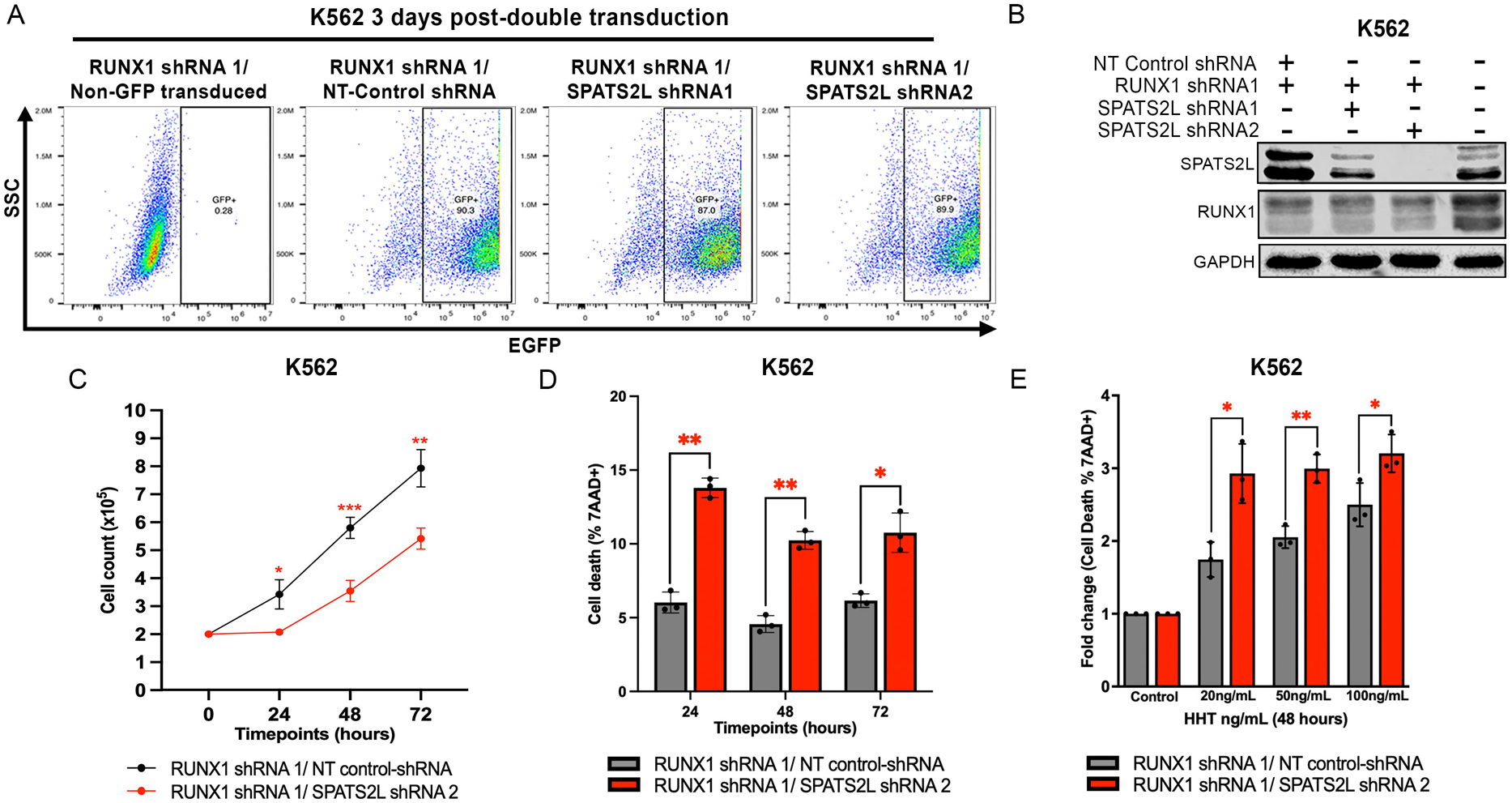
Inhibiting SPATS2L induction in RUNX1 depleted K562 cells impairs growth, viability and increases HHT sensitivity. **A)** Representative flow cytometric pseudoplots showing the percentage EGFP^+^ cells 3-days post double-transduction, in K562 RUNX1 shRNA 1 cells, transduced with two SPATS2L-targeting GFP^+^ shRNAs compared to a non-targeting GFP^+^ NT control-shRNA. These plots have also been shown alongside non-GFP^+^ shRNA transduced/untransduced RUNX1 shRNA 1 cells. GFP^+^ populations (rectangle) are annotated as a percentage of all cells analysed. **B)** Western blot showing level of SPATS2L and RUNX1 in K562 RUNX1 shRNA 1 cells, transduced with two GFP^+^ SPATS2L-targeting shRNAs, compared to a GFP^+^ non-targeting shRNA. GAPDH was used as a loading control. **C)** Line graph summarising cell growth rates of K562 RUNX1 shRNA 1 cells transduced with a GFP^+^ NT control-shRNA or SPATS2L shRNA 2 at 24- 48- and 72-hours. Data represents mean ± 1 s.d (*n = 3*) and statistical significance is denoted by *p<0.05, **p<0.01 or ***p<0.001 from a two-tailed unpaired student’s t-test. **D)** Summary bar graph showing viability of K562 RUNX1 shRNA1/NT control-shRNA and K562 RUNX1 shRNA1/SPATS2L shRNA2 cells as deduced by flow cytometric assessment of 7AAD^+^ (dead cells). Data represents mean ± 1 s.d (*n = 3*) and statistical significance is denoted by *p<0.05 or **p<0.01, from a two-tailed unpaired student’s t-test. **E)** Summary bar graph showing the fold change in percentage of dead cells in K562 RUNX1 shRNA1/NT control-shRNA cells compared with RUNX1 shRNA1/SPATS2L shRNA2 cells, following 48-hours HHT treatment (20-100ng/mL) as deduced by flow cytometric assessment of 7AAD^+^ events. Data from each cell line are normalised to the control dose, mean ± 1 s.d (*n = 3*). Statistical significance is denoted by *p<0.05 or **p<0.01 from a two-tailed unpaired student’s t-test on log_2_ transformed fold change values.

## Discussion

Using the RNA-interactome capture method, OOPS, we characterised how RUNX1 loss causes post-transcriptional dysregulation in BP-CML cells. RUNX1 is a regulator of post-transcriptional processes including splicing ^47–49^ and RiBi/translation.^36,42^ We found RUNX1 depletion in BP-CML cells impacted RBPs associated with stress responses and RiBi/translation. Previous research has shown different *RUNX1* mutations in the RHD cluster into LOF, hypomorphic and wild-type like groups;^50^ hypomorphic mutants are characterised by defective CBFβ interaction.^51^ LOF/hypomorphic *RUNX1* mutations are associated with the enrichment of genes with stress response pathways and rRNA processing/RiBi,^50^ supporting our findings seen in KU812 cells with RUNX1 depletion, implying the induction of the stress response and dysregulated RiBi/translation are intrinsically linked. Consequently, the interaction between these two processes led us to consider whether RUNX1, known to be a regulator of rRNA transcription,^36^ could function more broadly as a regulator of 5’TOP motifs. Cells exposed to stress downregulate cap-dependent translation through the phosphorylation of eIF2α, modifying the expression of mRNAs containing a 5’TOP motif.^38^ We found multiple 5’TOP motifs were dysregulated with RUNX1 depletion in BP-CML cells. Notably, constituents of ribosomal subunits were predominantly increased in their expression. We also found an increase in ribosome occupancy in polysomes with RUNX1 depletion, without an increase in actively translating polysomes, suggesting global translation does not increase. As cells adapt to stress, where translation is initially arrested, the modification of ribosome production/function occurs, facilitating translational reprogramming, critical to the cells survival.^52^ Thus, *RUNX1* mutations in CML could promote transformation through translational reprogramming following the induction of stress.

The disruption of cap-dependent translation leads to canonical SG-assembly; membraneless cytoplasmic foci which protect mRNAs/proteins from degradation, promoting survival and therapeutic resistance.^33,34^ We found RUNX1 depletion or expression of RUNX1::EVI1 induced SPATS2L, a component of SGs.^31,32^ In BP-CML cells, we found SPATS2L depletion inhibited growth, survival and reduced SG-assembly, in keeping with previous research.^32^ Supporting our findings, *SPATS2L* has been shown to be upregulated in *RUNX1* mutated, compared with *RUNX1* wild-type BP-CML cells.^23^ Furthermore, we found induction of *SPATS2L* mRNA in BP-CML cells with RUNX1 depletion or expression of RUNX1::EVI1. However, RUNX1 ChIP-peaks are not enriched at the promoter of *SPATS2L* consistently in different contexts,^53,54^ indicating the induction of *SPATS2L* with RUNX1 dysregulation could be through altered mRNA stability/decay in response to cellular stress. SPATS2L has been shown to regulate JAK/STAT ^40^ and type I interferon signalling,^55^ and, *RUNX1* mutations in BP-CML are associated with upregulated interferon signalling.^23^ This suggests SPATS2L could contribute to the activation of this pathway, by potentially regulating SG-assembly, which serve as critical signalling hubs in type I interferon responses.^56^ Notably, we found BP-CML cells with RUNX1 depletion had an increase in SG-assembly, suggesting these cellular components could be critical to *RUNX1*-mutant pathogenesis.

The induction of SG-assembly with RUNX1 depletion in BP-CML cells led us to consider whether their targeting represents a novel therapeutic strategy. HHT and its semisynthetic derivative Omacetaxine Mepessucinate (OM), bind to the ribosomal-A site preventing new ribosomes entering elongation, but allowing ribosomes already in elongation to run-off mRNAs.^57–59^ Harringtonine inhibits SG-assembly by freezing 80s ribosomes on mRNAs, changing their conformational structure and preventing the recruitment of ‘ribosome-free’ mRNAs into these condensates.^41,57^ We found HHT ablated SG-assembly in RUNX1 deficient K562 cells. Notably, it has been suggested BCR::ABL1 localises to SGs, essential for its oncogenic function,^60^ implying that the inhibition of SGs could also disrupt BCR::ABL1 function. HHT has a history of clinical use in CML ^43–46, 61^ and has demonstrated pre-clinical efficacy in *RUNX1* mutated AML cells.^42^ These findings were followed up with a phase I/II clinical trial where no responses were seen in relapse/refractory *RUNX1* mutated AML patients treated with OM/Venetoclax.^62^ However, doses used were lower than previously used in CML ^43–46^ and two patients with untreated *RUNX1* mutated MDS had complete composite responses, leading the authors to indicate that HHT/OM could be beneficial in non-heavily pre-treated *RUNX1* mutated myeloid malignancies.^62^ As CML represents a disease not heavily pre-treated before the advanced-phases, we considered whether BP-CML cells with *RUNX1* mutations could also be sensitised to HHT.

*RUNX1* mutations in BP-CML cells exhibit different sensitivities to novel therapies, highlighting their heterogeneity.^23^ On testing primary *RUNX1* mutated BP-CML cells with HHT, we found *RUNX1p.K117*, RUNX1p.R107C* and *RUNX1p.S100F*, existing within the RHD, exhibited the greatest sensitivity. *RUNX1p.K117\** (*RUNX1p.K90fsx101*- isoform b) causes haploinsufficiency through defective CBFβ interaction, nuclear localisation, DNA-binding and transactivation.^63^ *RUNX1p.R107C* (*RUNX1p.R80C-* isoform b) occurs at a ‘hotspot residue,’ leading to defective CBFβ interaction, DNA-binding and transactivation.^64^ Finally, *RUNX1p.S100F* (*RUNX1p.Ser73Phe-* isoform b) causes defective CBFβ interaction and transactivation.^65^ These findings imply that the preferential sensitisation to HHT of the *RUNX1* mutations we tested is due to the disruption of DNA-binding/CBFβ interaction. *RUNX1* RHD mutations clustering into LOF/hypomorphic groups, not wild-type like, are enriched for a gene signature associated with stress response pathways and RiBi.^50^ This further implies that the increased sensitivity to HHT we found with the LOF/hypomorphic *RUNX1* mutations we tested is related to the dysregulation of these post-transcriptional processes. Supporting these findings, the *RUNX1* C-terminal mutations we tested, including *RUNX1p.R320\** and *RUNX1p.Q274fs*His36*, were less sensitised to HHT than BP-CML cells with wild-type *RUNX1*. The *RUNX1p.R320\** mutation does not affect CBFβ interaction ^65^ and exhibits similar genomic binding sites to RUNX1,^66^ suggesting it drives leukaemogenesis through altered mechanisms. Furthermore, *RUNX1* mutations leading to C-terminal deletions bind to DNA with a higher-affinity than RUNX1, altering the transcriptional programme by affecting the activation/repression of target genes.^67^ Thus, the difference in response to HHT between LOF/hypomorphic-RHD and C-terminal mutations could be related to differences in the induction of dysregulated post-transcriptional processes, highlighting that a mutationally-targeted approach may be required.

LOF/hypomorphic *RUNX1* mutations are associated with the induction of stress response pathways, ER stress and PERK-regulated genes.^50^ The induction of the integrated stress response (ISR), through PERK-activation, leads to eIF2α phosphorylation,^68^ promoting SG-assembly. Furthermore, PERK-eIF2α phosphorylation is upregulated in CML cells which have become resistant to imatinib.^69^ These findings imply that SG induction could be a feature of CML cells with LOF/hypomorphic *RUNX1* mutations, promoting TKI-resistance. We found SPATS2L was induced with RUNX1 depletion or expression of RUNX1::EVI1. SPATS2L amplification occurs in AML cells with increasing resistance-indices to HHT.^40^ Thus, whilst the induction of stress response pathways, dysregulated RiBi/translation and SG-assembly, may sensitise BP-CML cells with LOF/hypomorphic *RUNX1* mutations to HHT, the continued amplification of SG-formation through a SPATS2L/SG axis could serve as a mechanism for increasing therapeutic resistance. To consider this, we created a K562 cell line with RUNX1 and SPATS2L depletion. Notably, K562 RUNX1 shRNA 1 cells, already sensitised to HHT, were further sensitised with concurrent SPATS2L depletion, suggesting SGs may contribute to therapeutic resistance. However, to have validated this we would need to evaluate SG-assembly dynamics in a K562 cell line with RUNX1 depletion with increasing resistance indices to HHT.

In summary, our study demonstrates RUNX1 to be a critical regulator of the RNA-bound proteome in BP-CML cells. We revealed the induction of stress response pathways, dysregulated RiBi/translation and increased SG-assembly, are a feature of BP-CML cells with *RUNX1* aberrations, and, LOF/hypomorphic *RUNX1* mutations may be preferentially sensitised to HHT through these dysregulated post-transcriptional processes. Whilst a limitation of our work was testing a small number of primary samples, due to the increasing rarity of BP-CML, our findings provide preliminary evidence in support of a mutationally-targeted approach in CML with *RUNX1* aberrations, and propose SPATS2L/SG targeting could represent a novel therapeutic approach in this disease.

## Supporting information

Polysome profiling AUC analysis

K562 OOPS MS data

KU812 IPA

KU812 OOPS MS data

## Acknowledgements

We thank University Hospitals Sussex (UHS) NHS Foundation Trust (Palmer PhD studentship) and the British Society of Haematology (BSH) for funding this work (Early-Stage Research Start-up Grant – Palmer). The authors also thank the patients involved for supplying samples for this research. Finally, the authors thank the staff at the Paul O’Gorman Leukaemia Centre (Glasgow) for generous help with FACS and the staff at Queen Mary University of London for generous help with OOPS. The visual abstract was created with BioRender.com.

## Author Contributions

DP devised the study, designed and executed experiments, analysed the data and wrote the manuscript. AM and RC assisted with primary BP-CML sample processing/analysis and PL, MW, KH performed proteomics analyses. GH, MC, SM, SA, KP provided BP-CML samples access and TC provided mentorship and scientific guidance. NB, SJ, and DF contributed experimental data whilst HW and MJW assisted with *RUNX1*::*MECOM/EVI1* cloning. FKM guided OOPS experiments and further scientific expertise were provided by SGK, AT and BPT. RGM co-devised the study, designed experiments, analysed data, managed the project and edited the manuscript.

