## Supplementary data for "*RUNX1* aberrations in blast-phase CML induce the RBP SPATS2L which promotes growth, survival and stress granule assembly"

### Supplementary methods, tables and results

### Supplementary methods

Cell lines/cell culture

K562 and KU812 cell lines were obtained from the Leibniz Institute DSMZ (DSMZ, Braunschweig, Germany). The SKH1 cell line was gifted from the Bonifer lab (University of Birmingham) gifted originally by Dr Mitani (Dokkyo Medical University, Japan). HEK-293T cells were gifted from the Tonks lab (University of Cardiff). Suspension cells were cultured in Roswell Park Memorial Institute-1640 media (RPMI-1640; Merck-Millipore, Dorset, UK), supplemented with 10% FBS (Biosera, Cholet, France), 100IU/mL penicillin/100µg/mL streptomycin (P/S; Merck-Millipore) and 2mM L-glutamine (Merk-Millipore), and HEK-293T cells in Dulbecco's Modified Eagle Medium (DMEM; Merck Millipore), with 10% FBS and penicillin/streptomycin (as above). Cell lines were maintained at ~2x10$^5$/mL with regular passaging, and kept at 37°C under a humidified atmosphere with 5% $CO_2$. For nuclear/cytoplasmic fractionations and immunofluorescence staining K562 and KU812 cells were treated with 0.5M sodium arsenite (Merck-Millipore) for 1-hour.

Cell lysates, protein quantification and western blotting

Approximately 1-5x10$^6$ cells were washed in cold phosphate buffered saline (PBS; Merck-Millipore) and centrifuged at 300 x g, 4°C, for 10 minutes. Cell pellet/s were resuspended in 50-100µL of 1X Cell Lysis Buffer (Cell Signalling Technology, Massachusetts, United States), containing 1.5mL of a 10X stock made up of, 20nM Tris-HCl [pH 7.5], 150mM NaCl, 1mM $Na_2$EDTA, 1mM EGTA, 1% Triton, 2.5mM sodium pyrophosphate, 1mM β-glycerophosphate, 1mM $Na_3VO_4$ and 1µg/mL leupeptin, diluted in 8.5mL de-ionised water, with the addition of a complete Mini Protease-Inhibitor Cocktail (PIC; Merck-Millipore). Cell lysates were kept on ice for 30 minutes, being vortexed every 10 minutes, before centrifugation for 10 minutes, at 17,000 x g, 4°C. The supernatant was removed and stored at -80°C. Protein quantification was performed using a detergent-compatible colorimetric assay kit (Bio-Rad, Hertfordshire, UK), using protein standards made up with bovine serum albumin (BSA; Thermo Fisher Scientific), diluted in de-ionised water and 1X Cell Lysis Buffer (range 6.25µg/mL-800µg/mL, with a control containing no BSA). Analysis of protein standards/cell lysates was carried out using the iMark Microplate Absorbance Reader (Bio-Rad) at 655nm wavelength through Microplate Manager (v6.3). Samples were diluted in 4X Laemmli sample buffer (Bio-Rad) and de-ionised water to give a final concentration of 1µg/µL and heated at 95°C for 5 minutes. PAGE was performed using gels of different acrylamide percentages (7.5-12.5%) (Severn Biotech, Kidderminster, UK), resolving/stacking buffer (Thermo Fisher Scientific), and de-ionised water, with the addition of ammonium persulphate (0.5g/mL)(APS; Thermo Fisher Scientific) and tetramethylethylenediamine (TEMED; Severn Biotech) to allow for solidification. Protein samples (range of 20-30µg) were loaded into the gels with 3-5µL of precision all blue protein standard ladder (Bio-Rad). Samples were electrophoresed (Bio-Rad) in running buffer (14.4g Glycine powder, 3g Tris; Thermo Fisher Scientific, 10mls 10% SDS solution; Severn Biotech, dissolved in 1L of de-ionised water) at 100V for 15 minutes, then 190V for up to 1 hour. Gels were removed and soaked in transfer buffer (14.4g Glycine powder, 3g Tris, 200mL methanol; Thermo Fisher Scientific, dissolved in 800mL of de-ionised water). Immobilon membranes (Merck-Millipore) were soaked in methanol for 5 minutes, then washed three times for 3 minutes in de-ionised water. The gel and immobilon membranes were compressed for 90 minutes in transfer buffer at 100V. The membrane was soaked in a 5% milk block, made by adding 2.5g milk powder to 50mL 1X Tris-buffered saline with Tween (TBST; 100mL 10X TBS; Severn Biotech, 1g Tween-20; Thermo Fisher Scientific, in 900mL de-ionised water), for 1 hour at room temperature under gentle agitation, before being soaked in primary antibodies (**Table S1**) overnight at 4°C with agitation. The following day membranes were washed in 1xTBST three times for 10 minutes, before a 1-hour incubation at room temperature with agitation in secondary antibodies (**Table S1**). Finally, membranes were washed in 1xTBST three times for 10 minutes. The LumiGLO chemiluminescence kit (Sera care, Milford, USA) was used for antibody bound proteins to be visualised using a Licor Odyssey (Image Studio software v5.2), being exposed for 10 minutes on the chemiluminescence channel with ladder detection at 700nm over 2 minutes.

<u>Preparation of plasmid DNA , Lentiviral preparation and transductions</u>

Competent cells harbouring lentiviral shRNA plasmids (**Table S2**), were streaked on to Luria Broth (LB; 10g of Bactotryptone, 5g of Bacto yeast extract; Thermo Fisher Scientific, and 10g of NaCl made up to 1L with molecular water) agar plates treated with 1µL ampicillin (Thermo Fisher Scientific) per 1mL of LB agar (1:1000). Colonies were grown overnight in an incubator at 37°C and single colonies picked using a sterile pipette tip, then cultured overnight in LB with the addition of 1µL ampicillin (Thermo Fisher Scientific) per 1mL LB (1:1000), being agitated at 260 rpm in a shaker at 37°C. Broths were centrifuged at room temperature, 2000 x g, for 15 minutes, before mini-preps using a GeneJET Plasmid Miniprep Kit (Thermo Fisher Scientific). Quantification of purified plasmid DNA was carried out using a NanoDrop 2000 spectrophotometer (NanoDrop 2000 v1.6.198; Thermo Fisher Scientific). For cell transfections 25cm$^3$ adherent cell culture flasks were coated in 2mls of poly-L-lysine solution (Merck-Millipore) for 30 minutes at room temperature, before it was replaced with 5mL of DMEM. HEK293T cells were washed in PBS, Trypsinised in 1mL, counted and seeded at a density of 5x10$^6$ per flask in 5mL DMEM, and cultured overnight. When they had reached >80% confluence, target plasmid DNA was transfected into HEK293T cells using a Lipofectamine™ 3000 kit (Thermo Fisher Scientific). For each flask, two separate sterile containers were prepared. In one container 666µL of OptiMEM media (Thermo Fisher Scientific), 2.2µg of envelope plasmid gifted from the Tonks lab (University of Cardiff) (pSL3pMD.2G/VSVOG), 4µg of packaging plasmid gifted from the Tonks lab (University of Cardiff) (psPAX2), 16µL of p3000 yellow cap reagent (Thermo Fisher Scientific), and 2.1µg of the plasmid DNA was added. In the second sterile container 675µL of OptiMEM and 19µL of p3000 red cap reagent was added (Thermo Fisher Scientific). These two containers were mixed together with gentle swirling, and incubated for 20 minutes at room temperature. The DMEM in each flask was reduced to 2mL and lipid-DNA complexes were added. Flasks were incubated overnight at 37°C, before the first harvest was taken off and replaced with 2.5mL of DMEM and incubated again overnight at 37°C. The next day, the second harvest was taken and combined with the first, followed by centrifugation at 300 x g for 5 minutes. Aliquots of 1mL were added to cryovials and snap frozen in liquid nitrogen before storage at -80°C. For lentiviral gene transductions, 40µg/mL retronectin (Takara Shiga, Japan) was dispensed into each of the required wells of a non-treated tissue culture 24-well plate (Thermo Fisher Scientific) and left overnight at 4°C. The next day the retronectin was removed and replaced with 1% BSA in PBS solution, and left for 30 minutes at room temperature. The lentivirus was collected from the -80°C freezer and thawed in a water bath at 37°C. The BSA solution was removed and replaced with 1mL of the relevant lentivirus. Plates were centrifuged at 2000 x g for 90 minutes at room temperature. After centrifugation, the supernatant was aspirated and replaced with cells at $2x10^5$/mL RPMI culture media. After 48-hours transduced cells underwent puromycin selection (1μg/mL) or flow cytometry/FACS to assess for $GFP^+$ expression.

Proteomic analysis of OOPS samples (University of Bristol Proteomics department)

In-gel digestion and TMT-labelling

Each experimental replicate per cell line condition/replicate containing 20μg of protein, was electrophoresed on a 10% SDS-PAGE gel, until the dye front had advanced by 1cm into the gel. Each gel lane was subsequently excised and subjected to in-gel tryptic digestion using a DigestPro automated digestion unit (Intavis Ltd.). The resulting peptides evaporated to dryness and were resuspended in 50μl of 100mM TEAB, labelled with Tandem Mass Tag (TMT) six-plex reagents according to the manufacturer's protocol (Thermo Fisher Scientific) and the labelled samples were pooled. Each pooled sample was then desalted using a SepPak cartridge according to the manufacturer's instructions (Waters, Milford, Massachusetts, USA). Eluate from the SepPak cartridge was evaporated to dryness and resuspended in buffer A (20 mM ammonium hydroxide, pH 10) prior to fractionation by high pH reversed-phase chromatography using an Ultimate 3000 liquid chromatography system (Thermo Fisher). Briefly, the sample was loaded onto an Xbridge BEH C18 Column (130Å, 3.5 μm, 2.1 mm X 150 mm, Waters, UK) in buffer A and peptides eluted with an increasing gradient of buffer B (20 mM Ammonium Hydroxide in acetonitrile, pH 10) from 0-95% over 60 minutes. The resulting fractions (concatenated into 4 in total) were evaporated to dryness and resuspended in 1% formic acid prior to analysis by nano-LC MSMS using an Orbitrap Fusion Lumos mass spectrometer (Thermo Scientific).

Nano-LC Mass Spectrometry

High pH RP fractions were fractionated using an Ultimate 3000 nano-LC system in line with an Orbitrap Fusion Lumos mass spectrometer (Thermo Scientific). Briefly, peptides in 1% (vol/vol) formic acid were injected onto an Acclaim PepMap C18 nano-trap column (Thermo Scientific). After washing with 0.5% (vol/vol) acetonitrile 0.1% (vol/vol) formic acid peptides were resolved on a 250 mm × 75μm Acclaim PepMap C18 reverse phase analytical column (Thermo Scientific) over a 150min organic gradient, using 7 gradient segments (1-6% solvent B over 1min, 6-15% B over 58min, 15-32%B over 58min, 32-40%B over 5min, 40-90%B over 1min, held at 90%B for 6min, and then reduced to 1%B over 1min with a flow rate of 300 nl $min^{-1}$. Solvent A was 0.1% formic acid and Solvent B was aqueous 80% acetonitrile in 0.1% formic acid. Peptides were ionized by nano-electrospray ionization at 2.0kV using a stainless-steel emitter with an internal diameter of 30μm (Thermo Scientific) and a capillary temperature of 300°C. All spectra were acquired using an Orbitrap Fusion Lumos mass spectrometer controlled by Xcalibur 3.0 software (Thermo Scientific) and operated in data-dependent acquisition mode using an SPS-MS3 workflow. FTMS1 spectra were collected at a resolution of 120 000, with an automatic gain control (AGC) target of 400 000 and a max injection time of 100ms. Precursors were filtered with an intensity threshold of 5000, according to charge state (to include charge states 2-7) and with monoisotopic peak determination set to Peptide. Previously interrogated precursors were excluded using a dynamic window (60s +/-10ppm). The MS2 precursors were isolated with a quadrupole Isolation window of 0.7m/z. ITMS2 spectra were collected with an AGC target of 10 000, max injection time of 70ms and CID collision energy of 35%. For FTMS3 analysis, the Orbitrap was operated at 30 000 resolutions, with an AGC target of 50 000 and a max injection time of 105ms. Precursors were fragmented by high energy collision dissociation (HCD) at a normalised collision energy of 60% to ensure maximal TMT reporter ion yield. Synchronous Precursor Selection (SPS) was enabled to include up to 10 MS2 fragment ions in the FTMS3 scan.

Data Analysis

The raw data files were processed and quantified using Proteome Discoverer software v2.4 (Thermo Scientific) and searched against the UniProt Human database (downloaded January 2024: 82415 entries) using the SEQUEST HT algorithm. Peptide precursor mass tolerance was set at 10ppm, and MS/MS tolerance was set at 0.6Da. Search criteria included oxidation of methionine (+15.995Da), acetylation of the protein N-terminus (+42.011Da) and Methionine loss plus acetylation of the protein N-terminus (-89.03Da) as variable modifications and carbamidomethylation of cysteine (+57.021Da) and the addition of the TMT mass tag (+229.163Da) to peptide N-termini and lysine as fixed modifications. Searches were performed with full tryptic digestion and a maximum of 2 missed cleavages were allowed. The reverse database search option was enabled and all data was filtered to satisfy false discovery rate (FDR) of 5%.

Bioinformatic analysis

For volcano plots of the RBP-interactome, comparison between each replicate was made using both raw and normalised data, and, the $\log_{10}$ p-value of each protein was plotted against the $\log_2$ fold change and visualised in R, using the ggplot2 package. Proteins identified where $p<0.05$ and $\log_2 FC>1$ or $p<0.05$ and $Log_2 FC<-1$ were highlighted. The downstream analysis of biological pathways/processes was conducted using Ingenuity Pathway Analysis (IPA; QIAGEN) based upon differentially expressed proteins ($p<0.05$) using the full dataset as a reference proteome. The heatmap was created using 'limma' in R/Bioconductor using a known list of 5'TOP motif proteins as a reference [1], using $Log_2$FCs comparing each condition to their respective control replicates.

RNA extraction/purification and One-Step RT-qPCR

RNA extraction/purification was carried out using the Quick-RNA™ Miniprep Kit (Zymo Research, USA). Between $1\text{-}2\times10^6$ cells were centrifuged at 500 x g for 3 minutes, and the supernatant was removed and each pellet resuspended in 300µL of RNA-lysis buffer and transferred to a Spin-Away™ filter column, then centrifuged at 14,000 x g for 1 minute. The flow-through was resuspended in an equal volume of 100% ethanol, before transferred to a Zymo-Spin™ IIICG Column. The column was subsequently centrifuged at 14,000 x g for 1 minute and the flow-through discarded. The column was washed in 400µL of RNA-wash buffer and centrifuged for 1 minute. The flow-through was discarded again and 400µL of RNA-prep buffer was added to the column before centrifugation again, at 14,000 x g for 1 minute. The flow-through was discarded and the column was washed twice more with 700µL and 400µL volumes of RNA-wash buffer, respectively, with centrifugation between washes (as above). The final flow-through was discarded and 100µL of DNase/RNase free water was used to elute the RNA from the column matrix following a final centrifugation at 14,000 x g for 1 minute. To remove genomic DNA contaminants from the RNA-sample the TURBO DNA-free kit™ (Invitrogen) was used. A total of 2µL of TURBO DNase (2U/µL) was added to each RNA-sample with 10µL of TURBO-DNase 10x buffer and each reaction was incubated at 37°C for 30 minutes. Following the incubation, 10µL of DNase Inactivation Reagent was added to each sample before vortexing and leaving at room temperature for 5 minutes. A final centrifugation at 10,000 x g was carried out for 5 minutes and the supernatant was removed and transferred to a new Eppendorf. Each RNA-sample was quantified using a NanoDrop spectrophotometer 2000 (Thermo Fisher Scientific) blanked against RNase free water. A One-Step SyGr RT-qPCR Kit (Apto-Life, London, UK) was used to detect the abundance of transcripts in the RNA-samples taken. The RT-qPCR reactions were prepared (**Table S3)** and conditions used for RT-qPCR listed (**Table S4)**. Forward and reverse primers used for RT-qPCR are listed (**Table S5)**.

Polysome profiling

A total of $10x10^6$ cells per experimental replicate were incubated at 37°C for 5 minutes with the addition of 100µg/mL of Cycloheximide (Merck-Millipore). They were centrifuged at 300 x g for 10 minutes, before each pellet was resuspended in 2mL of ice-cold PBS (Merck-Millipore) with the addition of 100µg/mL of Cycloheximide. They were centrifuged again (as above) and each pellet was resuspended in 500µL of lysing buffer made up of 10mM Tris-HCl [pH 7.5], 150mM NaCl, 10mM $MgCl_2$, 1% Triton X-100 (Thermo Fisher Scientific), 1% NP40 (Thermo Fisher Scientific), 2mM Dithiothreitol (DTT; Merck-Millipore), 100µg/mL Cycloheximide, 12U/mL TURBO DNase™ (Thermo Fisher Scientific), 200U/mL SUPERase·In™ (Thermo Fisher Scientific), 1x Mini Protease-Inhibitor Cocktail tablet (Merck-Millipore), 0.5% Sodium deoxycholate and RNase free water. The lysates were kept on ice for 30 minutes, with pipetting every 10 minutes. Each lysate was centrifuged for 10 minutes at 4,000 x g at 4°C. The supernatant was removed and centrifuged again for 10 minutes at 12,000 x g at 4°C. A total of 400µL of the supernatant for each replicate was loaded carefully onto a sucrose gradient prepared the day before. Sucrose gradients were made by combining a 15% and 60% sucrose solution made up in a final volume of 50mL using sucrose powder (Merck-Millipore), poured into centrifuge tubes (Beckman Coulter, Amersham, UK). Each 15% and 60% sucrose solutions were made up (**Table S6)**, and mixed using a BioComp GradientMaster machine and stored at 4°C until the next day. After loading lysates onto the pre-prepared sucrose gradients, they were ultracentrifuged using a SWT40I rotor with the following settings, Radius min (66.7mm), Radius max (158.5mm), RPM (31,000rpm), RCF average (121,355 x g), RCF max (170,920 x g), K. factor (228.6), Run time (3 hours 30 minutes), Run temperature (4°C). The polysome profile of each replicate was obtained using a Triax flow cell machine and analysed with Triax flow cell software (v2.40).

HHT-sensitivity testing and 7AAD/Annexin V staining

Homoharringtonine/HHT was purchased from SelleckChem, Houston, USA. For each cell line condition, the required number of cells were washed in 20mL PBS (Merck-Millipore) and centrifuged at 300 x g for 5 minutes. Each pellet was resuspended to give a density of $2x10^5$/mL in culture media in a 24-well tissue-culture plate (Thermo Fisher Scientific). Concentrations of 20, 50, 100, 200 and 500ng/mL HHT were used alongside a DMSO (Merck-Millipore) control matched to the highest volume of HHT. For 7AAD staining analysis was performed at 24-, 48- and 72-hour timepoints using flow cytometry. To do this ~$2x10^4$ cells (100µL) were removed from the culture, added to a 96 well v-plate, and centrifuged at 300 x g for 3 minutes. Residual media was removed and each cell pellet was resuspended in 150µL staining buffer (0.5% BSA in

PBS) containing 1μg/mL 7-aminoactinomycin D (7-AAD) viability staining solution (Invitrogen). To perform Annexin V staining the Annexin V Apoptosis Detection Kit (BD Biosciences, San Diego, USA) was used. To do this ~2x$10^4$ cells (100μL) were removed from the culture, added to a 96 well v-plate, and centrifuged at 300 x g for 3 minutes. Residual media was removed and each well was resuspended in 200μL of cold PBS, before being centrifuged again at 300 x g for 5 minutes. The residual PBS was removed and each pellet was resuspended in 150μL of 1X Annexin V binding buffer (made up in 10X sterile water), with a 1:10 concentration of Annexin V-FITC and 5μg/mL PI added to each well. The plate was incubated in the dark for 15 minutes at room temperature prior to analysis. Flow cytometric analyses for 7AAD and Annexin V staining, were performed using a Cytoflex (Beckman Coulter) in conjunction with CytExpert software (v2.6) and a minimum 10,000 debris excluded events were collected. Post-acquisition analysis was carried out using FlowJo software (v10.10) (BD Life Sciences, USA).

Preparation, testing and analysis of primary patient-derived BP-CML cells

A total of 4 *RUNX1* wildtype and 3 *RUNX1* mutated, patient-derived BP-CML samples were supplied from the MATCHPOINT trial biobank [2], which received UK Research Ethics Committee approval (13/SC/0583) carried out in compliance with the Declaration of Helsinki, and, 2 *RUNX1* mutated patient-derived BP-CML samples were supplied from the personalised medicine project, Helsinki, where all subjects gave written consent in accordance with the declaration of Helsinki [3]. Clinical/molecular/cytogenetic characteristics of these patients is listed (**Table S7)**. Each sample was thawed and transferred to its own 50mL Falcon tube (Merck-Millipore) dropwise over 20 minutes, with each tube containing 10mL of pre-warmed DAMP solution, made up of DNase I 5000U/2mL, $MgCl_2$ 1.0M 1.25mL, Trisodium citrate 0.155M 53mL (Merck-Millipore), 25mL of 20% human serum albumin, made up to a final volume of 500mL in PBS. Each sample was centrifuged at 230 x g for 10 minutes and the wash with 10mL DAMP solution was repeated twice. Each pellet was resuspended in 4mL of high growth factor media, made up of serum free media with the addition of 50μg/mL of SCF, FLT3, IL3, IL6, and GCSF (Thermo Fisher Scientific). Primary samples were incubated overnight at 37°C, 5% $CO_2$. The next day, each sample was pelleted in a 15mL Falcon tube at 230 x g for 10 minutes. The supernatant was removed and each pellet was washed in 5mL of 2% FBS/PBS. Each sample was subsequently pelleted again at 230 x g for 10 minutes and resuspended in 2mL of 2% FBS/PBS, before being transferred to a sterile FACS tube. Approximately 1x$10^5$ cells were removed for unstained/single stain colour controls. The remaining samples were pelleted again by centrifugation at 230 x g for 10 minutes, the supernatant was tipped off, and each sample was incubated in 0.4μg/mL Anti-Human Lineage Cocktail (FITC) and 0.3μg/mL Mouse Anti-Human CD34 (APC) in 100μL 2% FBS/PBS in the dark for 30 minutes (**Table S1**). Each sample was centrifuged at 230 x g for 10 minutes, before cell pellets were resuspended in a final volume of 500μL 2% FBS/PBS and kept on ice for FACS of $Lin^-CD34^+$ cells. Following FACS, each sample was centrifuged (as above), and pellets were resuspended in the required volume of low growth factor media- made up of high growth factor media diluted to 1:100 in SFM (Thermo Fisher Scientific). Based on viable cell numbers following FACS, cells were seeded at a density of ~1-5x$10^5$/mL per well. However, sample MP014 was plated at ~7x$10^3$ cells in 200μL over two-wells of a 96-well plate due to low viable cell numbers. Samples were treated with either 50ng/mL and/or 20ng/mL HHT, with each sample having a

DMSO control matched to the volume of 50ng/mL HHT. Experimental replicates were performed in duplicate where sufficient cell numbers allowed, and the average generated. At 48-hours the percentage of cells in early/late-stage apoptosis in each sample was analysed using Annexin V positive staining. Flow cytometric analyses were performed using a BD FACS Canto™ II Clinical Flow Cytometry System and a minimum 10,000 debris excluded events were collected. For sample MP014 only 2500 debris excluded events were collected. Post-acquisition analysis was carried out using FlowJo software (v10.10) (BD Life Sciences, USA).

Cloning protocol

The pME18s-RUNX1-EVI1 vector was originally gifted to the Bonifer lab (University of Birmingham) by Dr Mitani (Dokkyo Medical University, Japan). The pSIEW plasmid was gifted from the Bonifer lab (University of Birmingham). The *RUNX1-EVI1* gene was amplified from the pME18s-RUNX1-EVI1 vector with a 50μL PCR reaction using 25μg of template DNA, with a Phusion flash high fidelity PCR master mix (Thermo Fisher scientific). Forward and reverse primers incorporating BamHI (3') and AscI (5') sites were designed and sourced (**Table S8**). The amplification conditions used were 2 minutes at 95°C, followed by 30 seconds at 95°C, 30 seconds at 65°C, 65 seconds at 72°C, and 2 minutes at 72°C. Each cycle was repeated 30X. The PCR product was run on a 1% agarose gel, and purified with a Monarch® PCR purification kit (New England Biolabs; NEB, Hertfordshire, UK). The pSIEW lentiviral vector was digested at 37°C for 2 hours, with 30 units of AscI, 60 units of high-fidelity Bam-HI, and 10X cut smart buffer (NEB). The digested PCR product also underwent a restriction enzyme digest. The post-digestion pSIEW vector was treated with rAPid Alkaline Phosphatase (Roche, Hertfordshire, UK) for another 15 minutes at 37°C. The digested pSIEW vector and digested PCR products were run on 1% agarose gels (made up using 0.5g agarose powder in 50mL TAE buffer), confirming the luciferase had been cut from the pSIEW plasmid. The remaining vector and PCR product were cut out of gels and cleaned up with a gel extraction kit (QIAGEN), as per manufacturer guidelines. The digested *RUNX1-EVI1* PCR product and cut pSIEW vector were ligated together overnight on ice, which slowly melted to room temperature, with a T4 DNA ligase and 10X DNA ligase buffer (NEB), in a 1:1 insert to vector ratio. The ligated vector was used to transform One Shot Stbl3™ chemically competent E.coli cells (Thermo Fisher Scientific), which were spread onto ampicillin containing LB agar plates, as above. Individual colonies were picked and the *RUNX1-EVI1* gene insert was confirmed by PCR from each colony. PCR products were electrophoresed on a 1% agarose gel to confirm the *RUNX1-EVI1* gene insert was present, allowing colonies to be selected for mini-prep. Final purified plasmids were sent for whole plasmid nanopore sequencing (Full Circle labs, London), confirming the *RUNX1-EVI1* gene had been cloned into the pSIEW vector.

Nuclear/Cytoplasmic fractionations

Approximately 5x$10^6$ cells were washed in cold PBS (Merck-Millipore) and centrifuged at 300 x g for 10 mins, at 4°C. The pellets were resuspended in 250μL cytoplasmic lysis buffer (10 mM Tris-HCl [pH 7.5], 10mM NaCl, 1.5 mM $MgCl_2$ and 0.5% Igepal-CA630/NP40; Cell Signalling Technology, London, UK) including 1 tablet of Complete Mini Protease Inhibitor; Merck-Millipore) and kept on ice for 10 minutes, vortexed at 5 minutes. Samples were centrifuged at 800 x g for 5 minutes at 4°C, and the supernatant was taken off as the cytoplasmic fraction without disrupting the pellet. Remaining pellets were washed in 500μL of cold PBS before repeat centrifugation (as above). Centrifuged pellets were resuspended in 100μL nuclear lysis buffer (see cell lysates protocol), followed by 5 cycles of sonication (30 second bursts, with 30 second rest). Samples were kept on ice for 30 minutes with vortexing every 10 minutes. The nuclear lysate was centrifuged at 17000 x g for 10 minutes at 4°C, and the supernatant was taken off as the nuclear fraction. Both the cytoplasmic and nuclear fractions underwent protein quantification and western blotting.

Immunofluorescence staining and Confocal Laser Scanning Microscopy (CLSM)

Approximately $4x10^6$ cells were washed twice in PBS and centrifuged at 300 x g for 5 minutes. The residual cell pellet/s were resuspended in 1mL fixative (2%) paraformaldehyde (PF) solution in PBS and incubated for 20 minutes at room temperature with occasional agitation. This was followed by two washes with staining buffer (0.5% BSA in PBS) and centrifugation at 300 x g for 5 minutes. Pellet/s were resuspended in 1mL of quenching buffer (100mM glycine in PBS) followed by a further two washes in staining buffer and centrifugation (as above). Pellet/s were resuspended in 1mL permeabilization buffer (0.1% Triton TX100 in PBS) and incubated at room temperature for 5 minutes with agitation. A further two washes in staining buffer was performed with centrifugation (as above). Following this, cell pellets were resuspended in a total volume of 100μL of staining buffer containing 2μL (1:50 concentration) of G3BP1 (Alexa Fluor® 488 Conjugate) antibody or IgG Isotype (Alexa Fluor® 488 Conjugate) antibody (**Table S1**) and incubated for 30 minutes at room temperature in the dark. Sample were washed again twice in staining buffer and centrifuged (as above). The supernatant was removed and the pellet was resuspended in 1mL of staining buffer containing 1mg/ml 6-diamidino-2-phenylindole (DAPI) for 5 minutes with agitation. Each sample was washed again in staining buffer and centrifuged as above, before the supernatant was removed and the remaining cell pellet was resuspended in 1mL of staining buffer. Cells resuspended in staining buffer (0.5% BSA in PBS) were spread across 35mm confocal dishes (Avantor, Pennsylvania, USA). Confocal microscopy was carried out using a Zeiss LSM880 confocal microscope with a 40x/1.3 oil immersion objective. Images were created using the HeNe 633 nm laser for Alexa Fluor 488 (G3BP1/IgG isotype stain), and 405 nm laser for DAPI staining, using Zeiss ZEN software (Black edition v2.3).

Statistical analysis

Cell growth, viability and drug sensitivity assays

Statistical testing for cell counting, cell viability and HHT-sensitivity assays were performed using Prism v10 (Graphpad, USA). Unless otherwise stated, all experiments were performed in experimental triplicates with data presented as the mean ± 1 standard deviation (SD) ($n$ = 3). Statistical significance has been defined using a threshold of *$P$<0.05, **$P$<0.01, or ***$P$<0.001 from either a two-tailed unpaired/paired student's t-test as stipulated.

RT-qPCR

Analysis of RT-qPCR was performed using Prism v10 (Graphpad, USA). All experiments were performed in experimental triplicates with technical duplicate/triplicate readings taken per experimental replicate. RT-qPCR was represented by fold change on ΔΔCt readings, normalised against a GAPDH input.

Statistical significance was denoted as deduced from a two-tailed unpaired student's t-test based on ΔΔCt readings, defined using a threshold of $^{*}P<0.05$, $^{**}P<0.01$, or $^{***}P<0.001$.

Polysome profiling

Polysome profiles for each experimental replicate were analysed using Prism (v10), inputting the absorbance readings (260 nm) and aligning the profile for each replicate to the base of the monosome. 'AUC analysis' (Prism v10) was used to calculate the total peak area of either the monosome or polysome fractions (from the base of the monosome to the end of the polysomes). Fold change difference was used to analyse the Total peak area of either the 80s monosome or polysomes. The average of all fold change differences across experimental replicates per cell line condition was taken, to give a final fold change reading for either the monosome or polysomes, normalised to 1 in the control shRNA cell lines. Polysome/monosome ratios were calculated using the formular, Total peak AUC of polysomes / Total peak AUC of Monosome, to give the P/M ratio (<1/>1).

Stress Granule analysis

For the analysis of stress granules, a total of 10X stack images were taken per experimental replicate (~5-10 cells per stack). Images of each stack were analysed through FIJI (ImageJ; software version 2.12.0/1.54 series). To perform the analysis, brightness/contrast was adjusted for each image to remove background signal. A 'Z-project' was performed for each stack using 'Maximum intensity' settings. The image was converted to 'Binary' before particles were analysed using the 'Analyse particles' function. Any particle <0.2μm (micron) in size was excluded as a stress granule, as previously shown [4]. The total number of stress granules per stack/number of cells within that stack, was used to give the average number of stress granules per cell and the average size of stress granules in each stack was also calculated. The combined average for the 10X stack images was used for each experimental replicate. Data represents mean ± 1 SD ($n$ = 3) with statistical significance assessed using a two tailed unpaired student's t-test and significance defined using a threshold of $^{*}P<0.05$, $^{**}P<0.01$, or $^{***}P<0.001$.

### Supplementary methods Tables (S1-S8)

**Table S1**: Primary and Secondary antibodies for western blotting, flow assisted cell sorting (FACS) and immunofluorescence staining.

| Antibody | Species | Dilution | Manufacturer/number |
|---|---|---|---|
| **Primary antibodies** | | | |
| RUNX1/AML1 | Rabbit polyclonal | 1:1000 | Abcam- ab23980 |
| GAPDH | Mouse polyclonal | 1:1000 | Proteintech 10494-1-AP |
| EVI1 | Rabbit monoclonal | 1:1000 | Cell Signalling- C50E12 |
| AML1 anti-RHD | Rabbit polyclonal | 1:250 | Calbiochem- PC285 *Product discontinued* |
| SPATS2L | Rabbit polyclonal | 1:1000 | Proteintech – 16938-1-AP |
| phospho-eIF2$\alpha$ | Rabbit monoclonal | 1:500 | Cell Signalling #3398 |
| eIF2$\alpha$ | Rabbit polyclonal | 1:1000 | Cell Signalling #9722 |
| Lamin AC | Mouse monoclonal | 1:50,000 | Merck-Millipore |
| $\alpha$ Tubulin | Mouse monoclonal | 1:50,000 | Merck-Millipore |
| FastImmune™ Anti-Human Lineage Cocktail 1 FITC (lin 1) (CD3, CD14, CD16, CD19, CD20, CD56) | Mouse monoclonal | FACS of Lin- cells (see primary samples protocol) | BD Biosciences 340546 |
| APC Mouse Anti-Human CD34 | Mouse monoclonal | FACS of CD34+ cells (see primary samples protocol) | BD Biosciences 555824 |
| G3BP1 (E9G1M) (Alexa Fluor® 488 Conjugate) | Rabbit monoclonal | (immunofluorescence protocol) | Cell Signalling #94496 |
| IgG Isotype Control (DA1E) (Alexa Fluor® 488 Conjugate) | Rabbit monoclonal | (immunofluorescence protocol) | Cell Signalling #2975 |
| **Secondary antibodies** | | | |
| Anti-Rabbit IgG | Goat | 1:1000 | Merck-Millipore |
| Anti-Mouse IgG | Goat | 1:1000 | Merck-Millipore |

**Table S2**: Plasmids for lentiviral preparation.

| Plasmid reference/TRCN number | Plasmid identified in manuscript as: | Source |
|---|---|---|
| MISSION RUNX1 shRNA TRCN0000013659 | RUNX1 shRNA 1 | Merck Millipore |
| MISSION RUNX1 shRNA TRCN0000013660 | RUNX1 shRNA 2 | Merck Millipore |
| MISSION SPATS2L shRNA TRCN0000053099 | SPATS2L shRNA 1 | Merck Millipore |
| MISSION SPATS2L shRNA TRCN0000053100 | SPATS2L shRNA 2 | Merck Millipore |
| MISSION pLKO.1-puro Non-targeted shRNA control | Control shRNA | Merck Millipore |
| pLV[shRNA]EGFP-U6-hSPATS2L[shRNA3] | SPATS2L shRNA 1 $GFP^+$ | Vector-Builder (London, UK) |
| pLV[shRNA]EGFP-U6-hSPATS2L[shRNA4] | SPATS2L shRNA 2 $GFP^+$ | Vector-Builder (London, UK) |
| pLV[shRNA]-EGFPU6>Scramble-ID:VB230321-1431mhe) | Control shRNA $GFP^+$ | Vector-Builder (London, UK) |

**Table S3**: SyGr RT-qPCR master mix for RT positive and negative reactions.

| Reagent | +RT reaction (Volume) | -RT reaction (Volume) |
|---|---|---|
| 2X SyGr RT-qPCR Mix | 5µL | 5µL |
| 20X Reverse Transcriptase | 1µL | |
| Primers | Forward/Reverse Primer 0.25µL (each) | Forward/Reverse Primer 0.25µL (each) |
| Template (RNA) | 1µL (10ng/µL) | 1µL (10ng/µL) |
| RNase free water | 2.5µL | 3.5µL |

**Table S4**: Conditions used for the RT-qPCR reactions in the thermocycler.

| Cycle | Temperature | Time |
|---|---|---|
| 1 | 50°C | 10 minutes |
| 1 | 95°C | 1 minute |
| 40 | 95°C<br>60°C | 5 seconds<br>20 seconds |
| Melt | Instrument settings | |

**Table S5**: Forward and reverse primers for each target gene: Each primer pair was resuspended in the required volume of RNase free water to give a stock concentration of 100μM. The stock was then diluted (1:10) with RNase free water to give a 10μM primer concentration used for PCR/RTqPCR.

| Gene name | Forward | Reverse |
|---|---|---|
| *GAPDH* | 5'-ACAGTCAGCCGCATCTTCTT-3' | 5'-ACGACCAAATCCGTTGACTC-3' |
| *SPATS2L* | 5'AAGCAGCATCAAGCCAACAAA-3' | 5'TTCTCGCAGCCATTCATGGG-3' |
| *RUNX1-EVI1* | (**Table S8**) | |

**Table S6**: Reagents and concentrations/volumes required to make 15% and 60% sucrose gradients for polysome profiling.

| Reagent | Stock concentration | Volume/amount required for 15% or 60% sucrose gradient made up in a final volume of 50mL |
|---|---|---|
| Sucrose | | (for 15% solution 7.5g, for 60% solution 30g) |
| Tris-HCl [pH 7.6] | 1M | 2.5mL |
| $MgCl_2$ | 2M | 1.5mL |
| NaCl | 5M | 0.25mL |
| Cycloheximide | 10mg/mL | 500 μL |
| Protease inhibitor | 1 tablet dissolved in 1mL of RNase free water | 165 μL (of 1mL stock) |
| DTT | 1M | 50 μL |
| RNase free water | | Make up to a final volume of 50mL |

**Table S7**: Clinical/molecular/cytogenetic characteristics of primary patient-derived BP-CML samples used for HHT sensitivity testing. Information was populated using, [2] [3]. MP- relating to MATCHPOINT trial identifier, VUS - variant of uncertain significance, HR (Haematological Response), CR (Cytogenetic response), MR (Molecular response), MMR (Major Molecular Remission). *RUNX1* mutated samples are highlighted in grey.

| **Identifier/ Biobank no.** | **Disease status** | **BP-CML lineage** | ***RUNX1* mutation (ClinVar status)** | **Co-mutation/s (pathogenic/likely pathogenic listed)** | **Additional cytogenetic abnormalities (ACAs)** | **Clinical outcome** |
|---|---|---|---|---|---|---|
| **MP002** | Progression to BP-CML on imatinib | Myeloid | *RUNX1 wild-type* | *BCOR* | t(13;21)(q1;q2),+8 t(13;21)(q1;q2), +17, add(2)(q1), der(8), t(8;18)(q24;q1),der(18) | Deceased |
| **MP003** | Progression to BP-CML on bosutinib | Myeloid | *RUNX1p.S100F* (VUS) | *TET2* | t(9;22;22) | HR- complete, CR-partial, MR-nil. Deceased |
| **MP006** | Progression to BP-CML on imatinib | Myeloid | *RUNX1 wild-type* | *ASXL1, STAG2* | +8, +9 +10, +13, +14, +15, +19, +21, +21, +Ph | Deceased |
| **MP009** | *De novo* BP-CML | Myeloid | *RUNX1 wild-type* | *ASXL1* | None identified | HR-complete, CR-complete, MR-MMR. Allograft -> Alive (at trial end) |
| **MP011** | Progression to BP-CML on dasatanib (with 18 months prior treatment with imatinib) | Myeloid | *RUNX1 wild-type* | None identified | +Ph | CR-complete. Allograft->Alive (at trial end) |
| **MP014** | *De novo* BP-CML | Mixed | *RUNX1p.R320** (Pathogenic) | *TET2, KIT* | None identified | HR-complete, CR-complete, MR-MMR. Allograft -> Deceased |
| **MP017** | *De novo* BP-CML | Mixed | *RUNX1p.Q274fsH*36* (VUS) | *TET2, EZH2, ZRSR2* | None identified | HR-complete, CR-complete, MR-MMR. Allograft-> Alive (at trial end) |
| **See** [3] | *De novo* BP-CML | Myeloid | *RUNX1p.R107C* (Likely pathogenic) | *PHF6, for further mutations see* [3] | Not known | Alive at 10 months |
| **See** [3] | *De novo* BP-CML | Myeloid | *RUNX1p.K117** (Pathogenic) | *BCOR, BCORL1, for further mutations see* [3] | Not known | Alive at 6 months |

**Table S8**: Forward and reverse primers used for cloning *RUNX1-EVI1* into the pSIEW lentiviral plasmid. AscI and BamHI sequences are highlighted in bold on the forward and reverse primer sequences, respectively.

| Primer | Primer sequence | Tm° | Oligo no. |
|---|---|---|---|
| Forward | aattta**GGCGCGCC**aatgcgtatccccgtagatgc | 83.4 | 8821195098-10/0 (Merck-Millipore) |
| Reverse | tcgcat**GGATCC**tcatacgtggcttatggactg | 80.0 | 8821195098-20/0 (Merck-Millipore) |

### Supplementary methods references

[1] Cockman E, Anderson P, Ivanov P. TOP mRNPs: molecular mechanisms and principles of regulation. Biomolecules. 2020;10(7):969.

[2] Copland M, Slade D, McIlroy G, Horne G, Byrne JL et al. Ponatinib with fludarabine, cytarabine, idarubicin, and granulocyte colony-stimulating factor chemotherapy for patients with blast-phase chronic myeloid leukaemia (MATCHPOINT): a single-arm, multicentre, phase 1/2 trial. The Lancet Haematology. 2022; 9(2):e121-32.

[3] Adnan Awad S, Dufva O, Ianevski A, Ghimire B, Koski J, et al. RUNX1 mutations in blast-phase chronic myeloid leukemia associate with distinct phenotypes, transcriptional profiles, and drug responses. Leukaemia. 2021;35(4):1087-99, 2021.

[4] Cheng SJ, Gafaar T, Kuttiyatveetil JR, Sverzhinsky A, Chen C, et al. Regulation of stress granule maturation and dynamics by poly (ADP-ribose) interaction with PARP13. Nature communications. 2025;16(1):621.

### Supplementary results

### Figure S1

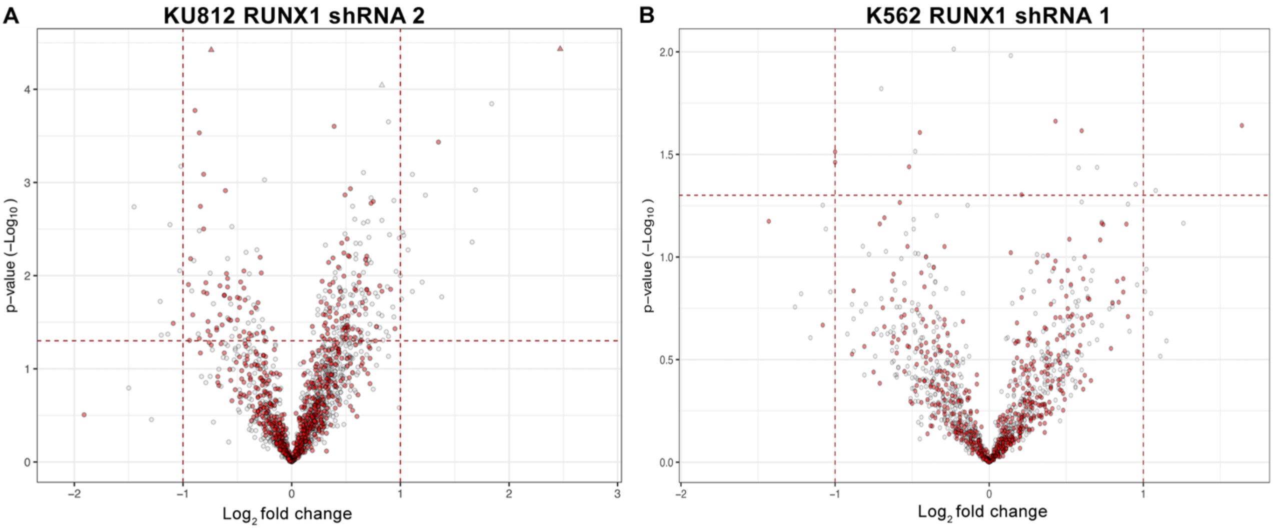


**Figure S1: The enrichment of RBPs from OOPS in RUNX1 depleted KU812 and K562 cells, A-B)** Volcano plots depicting the enrichment of RBPs in RUNX1 depleted KU812 (shRNA 2) and K562 (shRNA 1) cells. The enrichment of RBPs in the RNA-bound proteome fraction in KU812 RUNX1 shRNA 2 cells was 49% (872/1778) and K562 RUNX1 shRNA 1 cells 47% (565/1215). Horizontal dashed lines indicate the threshold for statistical significance ($p<0.05$) on a $\log_{10}$ scale (1.3) and vertical dashed lines represent $\text{Log}_2$FC (>1). Red dots represent RBPs (based on GO search term *RNA-binding*)and white dots are non-RNA-binding proteins.

### Figure S2

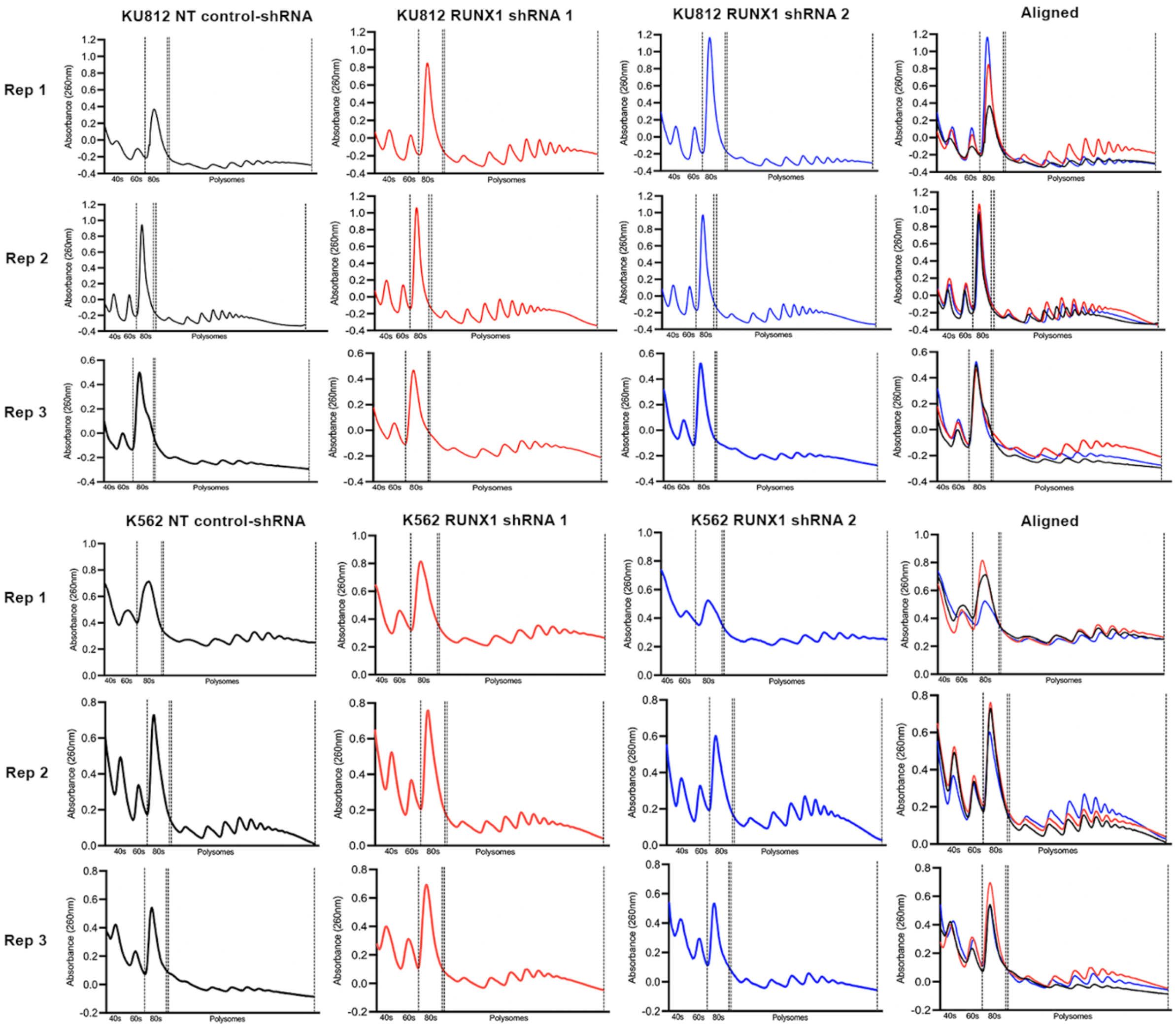


**Figure S2: Additional polysome profiling replicates.** Polysome profiles of KU812 and K562 cells (Replicates 1-3) transduced with NT control-shRNAs (black) compared to RUNX1-targeting shRNAs (red/blue). Dashed vertical lines represent monosome and polysome fractions. The 40s, 60s, 80s subunits, polysomes and absorbance readings (260nm) are annotated. Individual profiles are shown alongside aligned profiles for each replicate.

### Figure S3

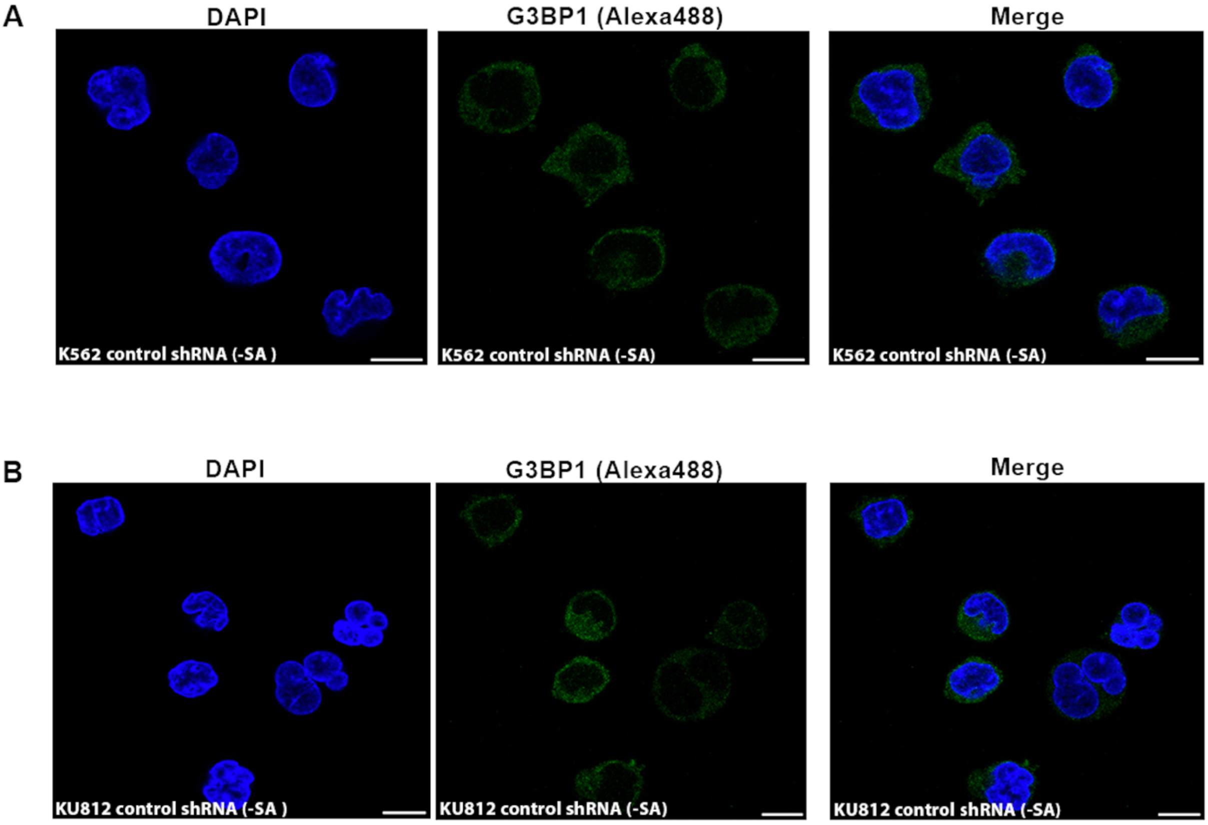


**Figure S3: Stress granules cannot be visualised with CLSM in K562 and KU812 NT control shRNA cells in the absence of sodium arsenite treatment (-SA). A-B)** Representative images show the DAPI, G3BP1(Alexa488), and merged stains, in K562 (A) and KU812 (B) control shRNA cells without sodium arsenite treatment (-SA). The scale bar in each image depicts 10 μm.

### Figure S4

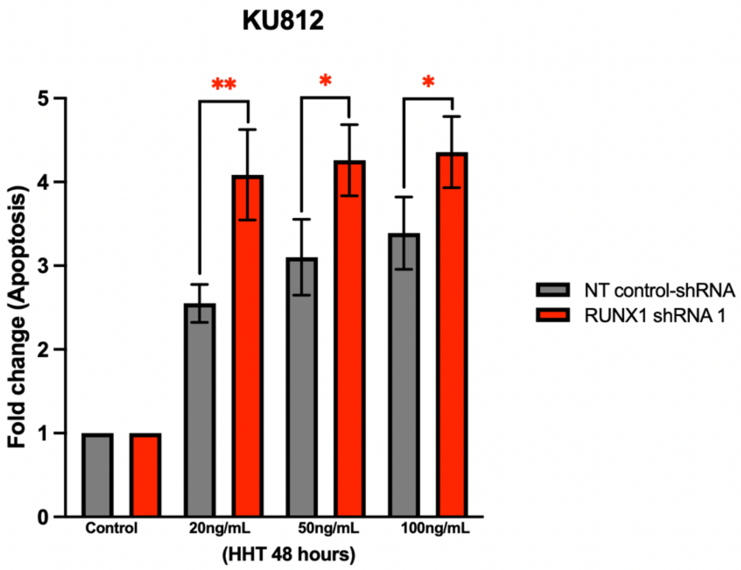


**Figure S4: KU812 RUNX1 shRNA 1 cells are sensitised to HHT through increased apoptosis.** Summary bar graph showing the fold change in percentage of apoptosis in RUNX1 shRNA1 compared to NT control-shRNA KU812 cells, following 48-hours HHT treatment (20-100ng/mL) as deduced by flow cytometric assessment of Annexin V positive (early/late-apoptosis) events. Data from each cell line are normalised to the control dose, mean ± 1 s.d (*n = 3*). Statistical significance is denoted by *p<0.05 or **p<0.01 from a two-tailed unpaired student's t-test on log2 transformed fold change values.
